# Generative Modeling of Mouse Embryogenesis for Fate and Disease Prediction

**DOI:** 10.64898/2026.06.18.733286

**Authors:** Yimin Fan, Xinyuan Liu, Yixuan Wang, Zehua Zeng, Lei Li, Xiaojie Qiu, Yu Li

## Abstract

Embryonic development is orchestrated by complex gene regulatory networks, and learning regulatory dynamics from developmental data could allow us to understand, predict, and ultimately engineer cell fates. Here we introduce Navigo (https://github.com/aristoteleo/Navigo-release), a biologically grounded generative modeling framework that learns a developmental vector field by integrating flow matching at the population level with RNA kinetics modeling at the molecular level. Navigo accurately maps developmental trajectories across lineages on a mouse embryogenesis scRNA-seq atlas spanning 43 time points and comprising 12.4 million cells. Applied to cardiac development, Navigo enables disease modeling by mechanistically resolving regulatory networks that distinguish congenital heart disease subtypes. Navigo also predicts perturbation effects in a zero-shot manner, as validated on independent *in vivo* data from six knockout genotypes without perturbation-specific training, uncovering lineage-specific gene-compensation mechanisms. Moreover, Navigo guides rational cell-fate engineering, exemplified by fibroblast reprogramming analyses, including identifying pro-fibrotic barriers to cardiac fates and evaluating hundreds of pairwise transcription factor combinations for neuronal fate, each consisting of one bHLH factor and one POU factor. Overall, Navigo provides a generalizable AI platform for perturbation-effect prediction, disease modeling, and rational cell-fate engineering, advancing toward AI-based virtual embryos for developmental biology and regenerative medicine.

## Introduction

Embryonic development represents one of nature’s most remarkable transformations, where a single fertilized cell orchestrates a precisely choreographed sequence of cellular decisions to generate the extraordinary diversity and complexity of a multicellular organism. This process is governed by gene regulatory networks (GRNs) operating through molecular interactions between regulators and their target genes, which collectively coordinate cell fate decisions, tissue morphogenesis, and organ formation [1, 2, 3]. Deciphering the regulatory principles underlying normal development addresses fundamental questions in developmental biology [4, 5], provides the foundation for interpreting how regulatory disruptions lead to congenital diseases [6, 7, 8], predicting the effects of genetic and regulatory perturbations, and rationally manipulating developmental programs to engineer desired cellular outcomes for regenerative medicine [9, 10]. Together, these goals motivate the emerging vision of a predictive virtual embryo: a generalizable *in silico* system that can be queried and perturbed to understand developmental mechanisms, predict disease-associated perturbation outcomes, and guide cell-fate engineering [11, 12].

Recent advances in single-cell RNA sequencing (scRNA-seq) have generated unprecedented temporal atlases of embryonic development [13, 14, 15, 16], profiling millions of cells across dozens of developmental time points from gastrulation through organogenesis to birth and capturing cellular states across diverse lineages and tissue types (Figure 1a,b). However, these snapshots capture only the outcomes of development at sampled time points. They reveal what cellular states exist at each measured moment but do not reveal how cells transition between states, what regulatory mechanisms drive these transitions, or how the system would respond to perturbations. Addressing these questions requires learning the underlying generative process itself: a continuous dynamical system that captures how regulatory networks drive cellular state evolution over time, encodes the regulatory logic governing lineage decisions, and enables counterfactual simulation of developmental outcomes under unobserved perturbations. Generative modeling [17, 18] provides this capability by capturing the data-generating process, transforming static observations into a dynamic, predictive, and programmable framework for modeling embryogenesis (Figure 1c).

**Figure 1.**
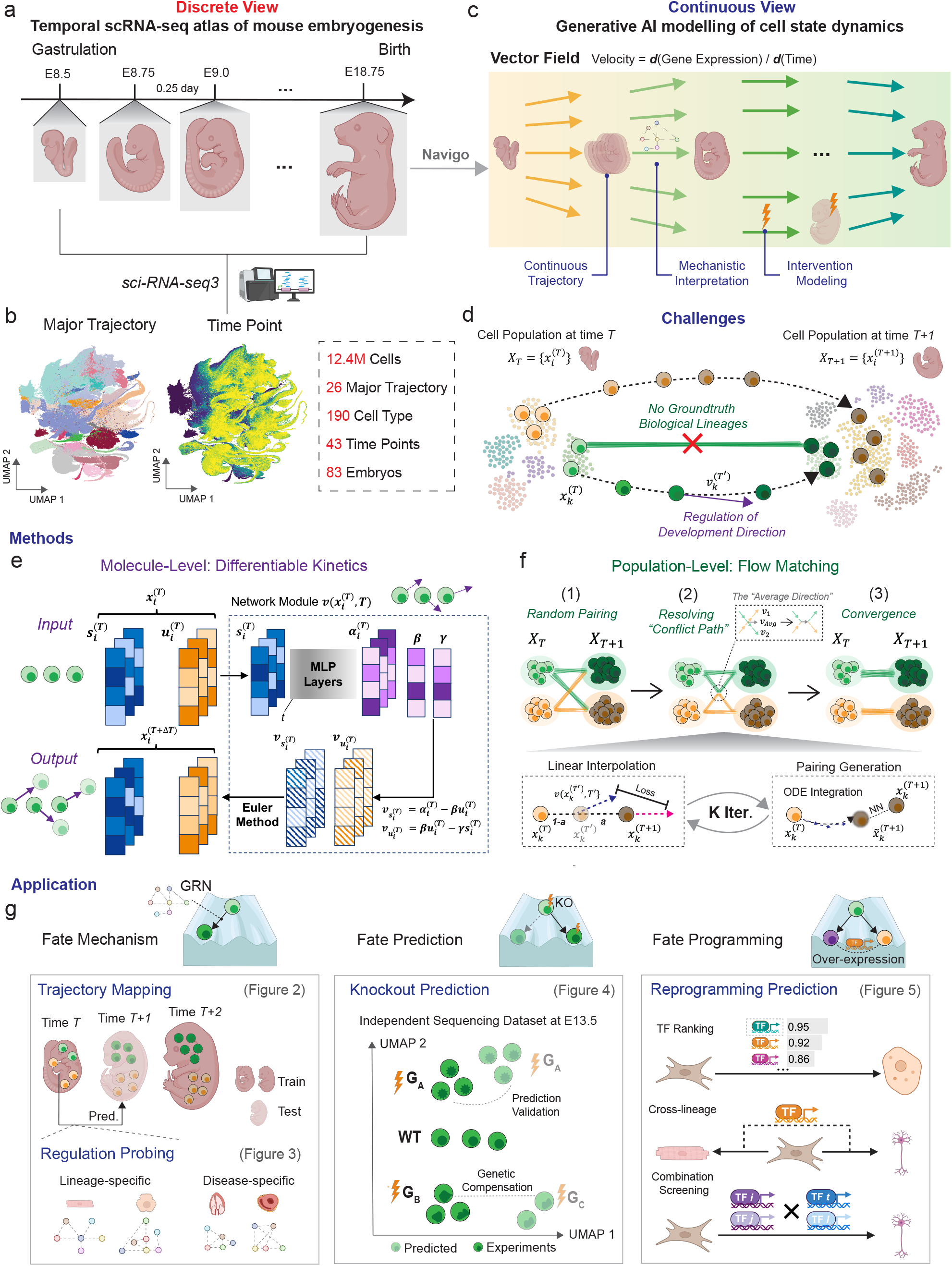
Overview of the Navigo Framework. **a.** Overview of the mouse embryogenesis atlas profiling developmental stages spanning from gastrulation (E8.5) to birth (E18.75) at 0.25-day intervals. The dataset was profiled using sci-RNA-seq3 technology, providing the most comprehensive static snapshots of the mouse embryogenesis process so far [13]. **b**. UMAP visualization of the mouse embryogenesis temporal atlas colored by major developmental trajectories (left) and time points (right), with accompanying dataset statistics. Detailed colormaps can be found in the Supplementary Figure 1. **c**. Navigo employs generative AI modeling to transform static snapshots into a continuous dynamic model of a vector field. The generative modeling framework enables three key capabilities that cannot be achieved from static snapshots alone: continuous trajectory inference across developmental time, mechanistic interpretation of gene regulatory dynamics, and *in silico* intervention forecasts. **d**. Key challenges for computationally modeling the embryogenesis process include inferring regulatory mechanisms governing developmental direction and the absence of ground-truth biological lineages. **e**. Model architecture. At the molecular level, Navigo integrates differentiable kinetics into a biologically grounded architecture. The model predicts transcription (*α*), splicing (*β*), and degradation (*γ*) rates based on spliced and unspliced gene counts, from which developmental velocities are inferred. **f**. Optimization objective. At the population level, the model is optimized using the rectified flow matching objective. The training procedure iteratively alternates between linear interpolation along paired cells across time, and pairing generation via ODE integration and nearest-neighbor matching, progressively refining from random pairings to convergent optimal trajectory. **g**. Key applications enabled by Navigo include fate mechanism understanding exemplified by trajectory mapping and regulatory network probing, fate prediction exemplified by zero-shot knockout prediction, and fate programming exemplified by reprogramming transcription factor prediction.

Current computational approaches for modeling developmental processes face distinct limitations. Early trajectory inference methods, including Monocle [19, 20] and Diffusion Pseudotime [21], reconstruct pseudotemporal ordering and branching structures from single-cell transcriptomes but do not explicitly model the underlying cellular dynamics. Methods based on RNA velocity [22, 23] capture molecular-level dynamics but do not explicitly model temporal information, while trajectory mapping [24] methods map population-level patterns but lack biological grounding in the molecular mechanisms driving transitions, limiting their ability to provide mechanistic insight. Neural-ODE-based approaches [25] can model continuous dynamics, but they can be computationally intensive and are not directly constrained by transcriptional kinetics. To address these gaps, we introduce Navigo, a biologically grounded generative modeling framework that learns developmental vector fields by integrating flow matching at the population level with RNA kinetics modeling at the molecular level. At the population level, Navigo employs flow matching [26] to efficiently learn continuous vector fields from discrete temporal snapshots, enabling the model to capture smooth developmental trajectories across the entire cellular state space. At the molecular level, Navigo explicitly models transcriptional kinetics [27] through differential equations governing RNA transcription, splicing, and degradation, grounding the learned dynamics in the mechanistic processes that drive gene expression changes. This integration enables Navigo to capture both the temporal trajectories of development and the molecular mechanisms underlying cell state transitions, thereby providing a scalable and generalizable foundation for disease modeling, perturbation-effect prediction, and rational cell-fate engineering.

We validate Navigo on a mouse embryogenesis scRNA-seq atlas [13] spanning embryonic day 8.5 (E8.5) to birth and comprising 12.4 million cells across 43 time points and 26 major trajectories (Figure 1b). Navigo accurately maps developmental trajectories across lineages, achieving superior performance in interpolating cellular states between observed time points. The learned generative model can also denoise cell populations affected by technical noise or sampling limitations by leveraging the continuous dynamics learned from population-level data. The power of this generative framework is demonstrated through three progressively complex biological applications spanning cell-fate mechanism understanding, prediction, and programming. First, in cell-fate mechanism understanding and disease modeling, we characterize regulatory mechanisms distinguishing congenital heart disease (CHD) subtypes by simulating perturbations of CHD-associated genes [28] and analyzing their transcriptomic responses, revealing distinct functional modules and disease-specific pathway signatures. Second, in cell-fate prediction, Navigo achieves zero-shot perturbation-effect prediction of six genetic knockout phenotypes on an independent E13.5 dataset [29] without perturbation-specific training. Further analysis of knockout responses within gene families reveals lineage-specific genetic compensation mechanisms, where the magnitude of the experimental knockout response depends on the predicted regulatory contribution of the perturbed gene relative to other family members. Third, in cell-fate programming, Navigo guides rational screening of transcription factor combinations for fibroblast reprogramming, successfully identifying established factors, discovering cross-lineage regulators [30], and achieving high AUROC in predicting hundreds of bHLH-POU TF combinations [31] for neuronal conversion in a zero-shot manner. Together, these results demonstrate that learning the generative process underlying development transforms static observational data into a generalizable AI platform for cell-fate mechanism understanding, prediction, and programming through *in silico* experimentation. By enabling disease modeling, perturbation-effect prediction, and rational cell-fate engineering within a single generative framework, Navigo advances toward AI-based predictive virtual embryos for developmental biology and regenerative medicine.

## Results

### Navigo Learns Dynamic Vector Fields from Temporal Atlas of Mouse Embryogenesis via Generative Modeling

Modeling embryogenesis from single-cell temporal atlases poses two fundamental challenges (Figure 1d). First, embryogenesis and developmental transitions are governed by high-dimensional non-linear regulatory mechanisms that are infeasible to measure directly given the current technologies. Models must therefore learn biologically meaningful transition dynamics driven by underlying gene regulatory networks (GRNs), rather than merely fitting arbitrary mathematical transformations between cell state distributions at discrete time points. Second, the destructive nature of single-cell sequencing eliminates ground-truth cell-to-cell correspondences across time points, making it impossible to directly observe individual cellular trajectories or lineage relationships. This requires models to infer developmental paths from population-level distributions without access to true cell pairings.

To address these challenges, Navigo employs a generative modeling framework that learns the continuous developmental vector field from temporal snapshots through joint modeling of molecular and population dynamics. At the molecular level, Navigo captures transcriptional dynamics using differentiable RNA velocity models (Figure 1e). The framework predicts biologically interpretable kinetic parameters from smoothed unspliced and spliced RNA counts: transcription rates (***α***), splicing rates (***β***), and degradation rates (***γ***). Transcription rates are modeled as functions of mature mRNA abundance, reflecting the principle that gene expression is regulated by upstream regulatory proteins, for which mature mRNA counts are used as a proxy for protein abundance [32, 33]. This design makes transcription rates both cell-specific and gene-specific, capturing cell-state dependent regulatory dynamics, while splicing and degradation rates remain gene-specific parameters, following previous studies [23, 32]. These kinetic parameters then define the velocity field through RNA kinetic equations, enabling the model to learn cell-specific regulatory dynamics that drive developmental transitions through a biologically grounded biophysical model of RNA transcription, splicing and degradation.

At the population level, Navigo addresses the absence of ground-truth correspondences by inferring transport paths between cell state distributions across time points using rectified flow matching with iterative reflow (Figure 1f). Initial random pairings between cells at consecutive time points create a coupling structure with many crossing connections. Navigo then iteratively alternates between training the vector field to match velocities along the current paired paths and updating the pairings via ODE integration under the learned field. Across reflow iterations, the coupling structure is progressively “unwired,” transforming from tangled crossing paths into straighter, more coherent trajectories. This iterative refinement allows the model to infer plausible developmental paths without requiring prior cell correspondence information. To improve scalability, Navigo employs metacell aggregation, which reduces dataset size by grouping transcriptionally similar cells while preserving rare cell populations to maintain biological diversity (*Methods*). This integrated approach enables Navigo to capture both temporal progression and molecular mechanisms, supporting various applications (Figure 1g), including fate mechanism understanding (Figure 2,3), fate prediction (Figure 4), and fate programming (Figure 5).

**Figure 2.**
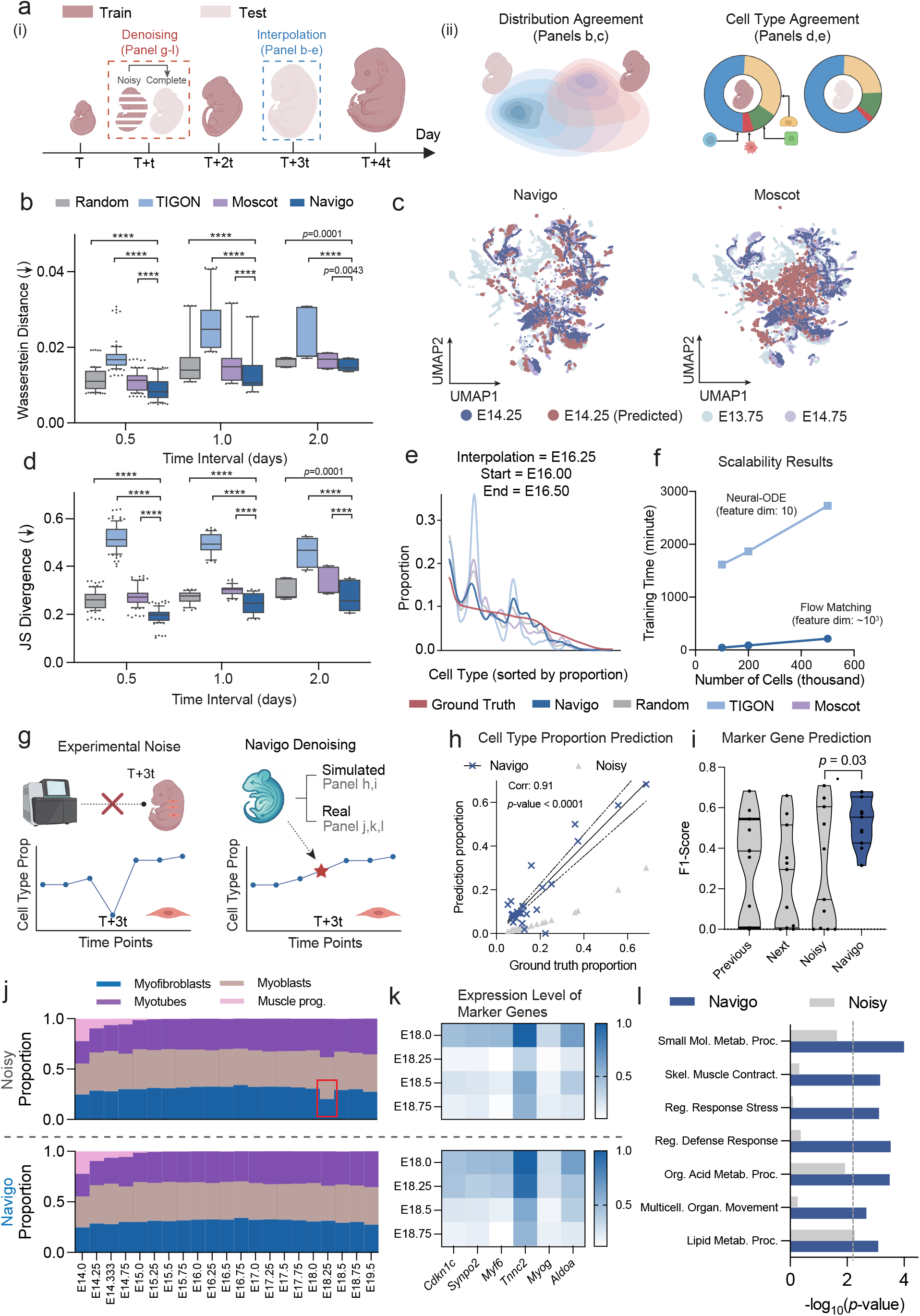
Navigo accurately maps and denoises the mouse embryogenesis trajectory on a comprehensive atlas. **a.** Overview of the study design: (i) schematic illustration of the interpolation and denoising tasks; (ii) evaluation of model performance using two metrics, distributional concordance and cell type agreement. **b**. Wasserstein distance comparison across different time intervals for distributional concordance between Navigo, Moscot, TIGON, and naive baseline methods. Time interval denotes the temporal gap between consecutive training time points. Box plots display median (center line), interquartile range (box, 25th–75th percentiles), and whiskers (10th–90th percentiles). Statistical significance was assessed using two-sided Wilcoxon matched-pairs signed-rank tests. Statistical significance is denoted by asterisks (four asterisks denote *p* ≤ 0.0001). *n*=95,40,15 for interval=0.5,1,2. **c**. UMAP visualization comparing interpolation predictions from Navigo and Moscot. Three time points are shown: start (E13.75), end (E14.75), and interpolated (E14.25). **d**. Jensen-Shannon (JS) divergence comparison for cell type agreement between Navigo and baseline methods across three time intervals. Box plot configuration and statistical analysis follow the same convention as in the panel **b. e**. Predicted cell type frequency distributions from Navigo and baseline methods across cell types on the x-axis, ordered by ground truth frequency. **f**. Computational scalability evaluation with respect to varying cell numbers. Panels **e** and **f** share the same legend. **g**. Schematic illustration of Navigo’s denoising capability. Denoising addresses challenges in experimental data where large cell populations may be inadequately captured due to technical noise or sampling limitations, evaluated on both simulated and real experimental datasets. **h**. Evaluation of predicted cell type proportions in simulated datasets. Cell type-time pairs with fewer than 10 cells were filtered for stability. Dashed lines indicate the 95% confidence interval around the regression line. “Noisy” denotes the simulated dataset with added noise. Association was assessed using Pearson correlation. *n* = 27. **i**. Concordance of differentially expressed genes (DEGs) between predicted and ground-truth cells of the same cell type in simulated datasets. For each cell type, overlap between the two DEG sets was quantified by the F1-score. Statistical significance was assessed using two-sided Wilcoxon matched-pairs signed-rank tests. *n* = 11. **j**. Comparison of original and Navigo-predicted cell type proportions along the muscle cell developmental trajectory across time points. Myofibroblast proportions at E18.25 exhibit apparent noise in the original data. **k**. Marker gene expression dynamics along the myofibroblast developmental trajectory in original versus predicted cell populations. **l**. Enriched biological pathways in original versus predicted myofibroblast populations at E18.25. We used the hypergeometric test to calculate the *p*-value.

**Figure 3.**
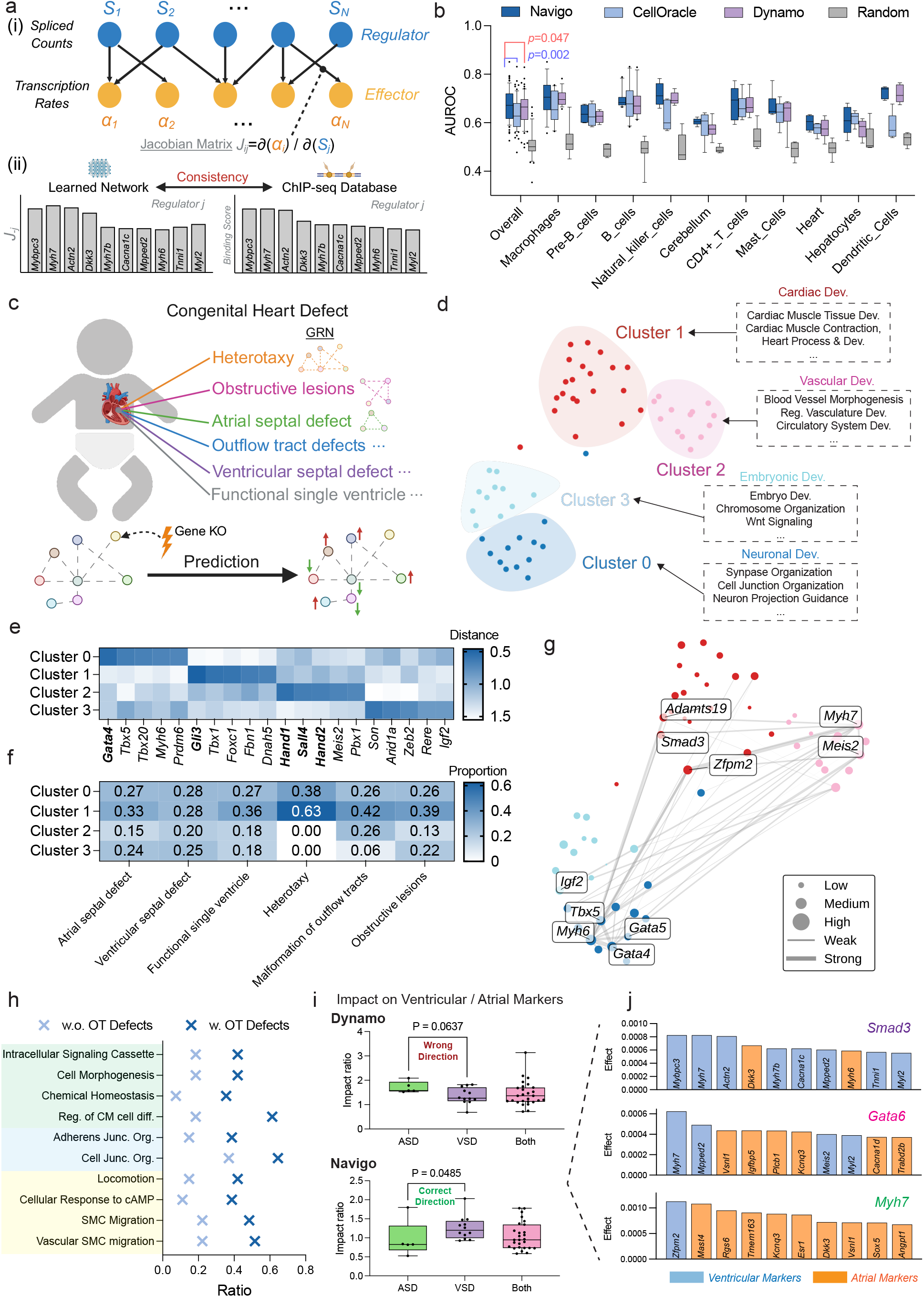
Navigo learns complex gene regulatory networks without prior knowledge. **a.** Illustration of the Jacobian network learned from the Navigo framework (i) and the evaluation procedure (ii). Networks are benchmarked for consistency with ChIP-seq databases across developmental lineages. **b**. AUROC scores comparing the network inferred from different methods against that of ChIP-seq databases across developmental lineages. We used two-sided Wilcoxon matched-pairs signed-rank tests to assess statistical significance (*n*=67). **c**. Schematic of CHD GRN characterization. Different CHD subtypes are driven by distinct GRNs. Navigo quantifies subtype-specific GRNs by simulating knockout and measuring transcriptomic responses. **d**. UMAP visualization of gene knockout response vectors averaged across cells reveals functional clustering. Each point represents the average transcriptomic response across cells to knocking out a specific CHD-associated gene. Clusters correspond to distinct developmental processes. **e**. Top CHD genes in each cluster ranked by minimum distance to the cluster center. Bold genes highlight the five genes with the smallest average distance across clusters. **f**. Proportion of CHD subtype-associated genes in each response-vector cluster, with proportions summing to 1 for each subtype. **g**. Regulatory interaction network of CHD genes. Edge thickness indicates mutual regulatory strength between genes. **h**. Pathway enrichment analysis of knockout-induced transcriptomic responses, stratified by outflow tract (OT) defect association. For each gene, pathway enrichment was computed from the gene knockout response vector. Ratio denotes the proportion of genes in each OT defect group whose knockout response was significantly enriched for a given pathway (hypergeometric test, *p <* 0.05). **i**. Impact of ASD- and VSD-associated genes on ventricular and atrial marker expression. Ventricular and atrial markers were defined as the top 50 upregulated differentially expressed genes for each cell type. *p*-values were calculated using a two-sided Wilcoxon rank-sum test. Gene numbers: ASD (5), VSD (12), Both (27). **j**. Examples of defect-specific regulatory effects. Top 10 regulated genes by *Smad3* (ASD-specific), *Gata6* (VSD-specific), and *Myh7* (shared). Colors indicate ventricular (blue) or atrial (orange) identity.

**Figure 4.**
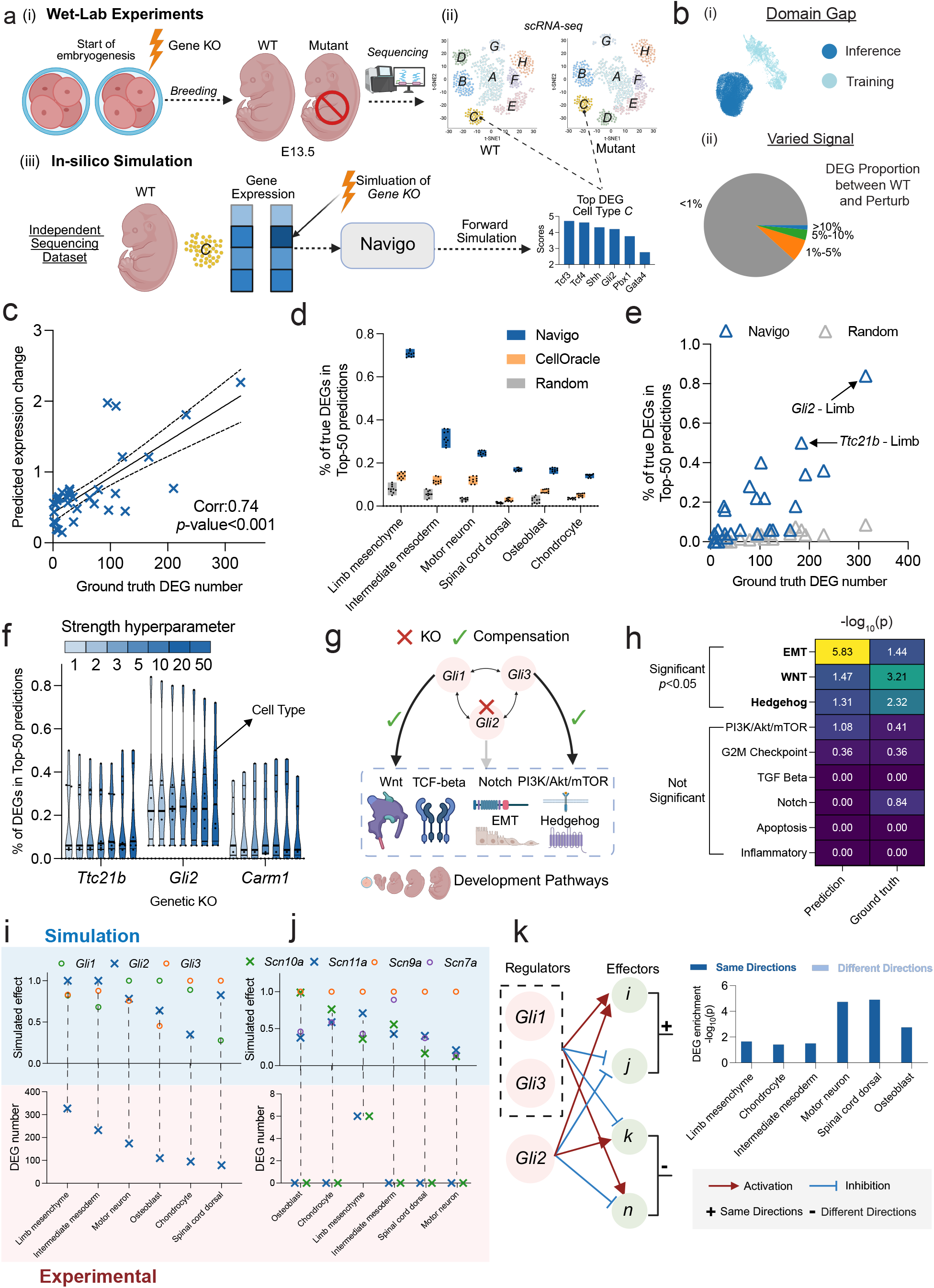
Navigo predicts genetic perturbation outcomes without training on specific knockout datasets. **a.** Overview of Navigo’s inference of genetic knockout effects during mouse embryogenesis using large-scale scRNA-seq data. (i) Knockouts are introduced at embryogenesis onset, and embryos are collected at E13.5 for scRNA-seq profiling of wild type and mutant samples. (ii) Top affected genes for each knockout and cell type are identified by differential expression (DE) analysis. (iii) Navigo simulates developmental trajectories and predicts the most affected DE genes, enabling direct comparison with experimental results. **b**. Prediction challenges: (i) the inference dataset is from an independent scRNA-seq study, resulting in substantial batch effects relative to the training data; (ii) for most gene knockout-cell type combinations, the proportion of DEGs between wild type and perturbation is low, as not every knockout produces significant changes in all cell types. **c**. Pearson correlation between the number of ground truth DEGs and absolute predicted expression changes across different gene KO and cell type pairs (*n*=35). **d**. Relationship between knockout effect size, defined by the number of ground truth DEGs, and prediction accuracy. **e**. Performance on the *Gli2* KO effect prediction on different cell types. **f**. Assessing the robustness of the predicted results in terms of the perturbation strength hyperparameter. **g**. Illustration of the gene compensation process. In the case of *Gli2* KO, genes in the same family could compensate for the effects of *Gli2* KO, thereby maintaining the function of development pathways. **h**. Comparison between the enriched pathways of the predicted top DEGs and the ground truth DEGs. **i**. Comparison of the simulated effects of the genes in the Gli family and the number of DEGs across different cell types. The simulated effect is quantified by the gene expression change after simulated KO and normalized by the maximal effect in the family. The DEG number is the ground truth DEG number of *Gli2* KO. **j**. similar to panel i, but for the *Scn* family. The DEG number is the ground truth DEG number of *Scn10a*/*11a* KO. **k** Regulatory directionality predicts gene sensitivity to *Gli2* knockout. Left: Predicted *Gli* effectors grouped by comparing simulated *Gli1*/*Gli3* double KO with *Gli2* KO directions. Right: Enrichment of experimental *Gli2* KO DEGs in “Same Directions” versus “Different Directions” categories across lineages.

**Figure 5.**
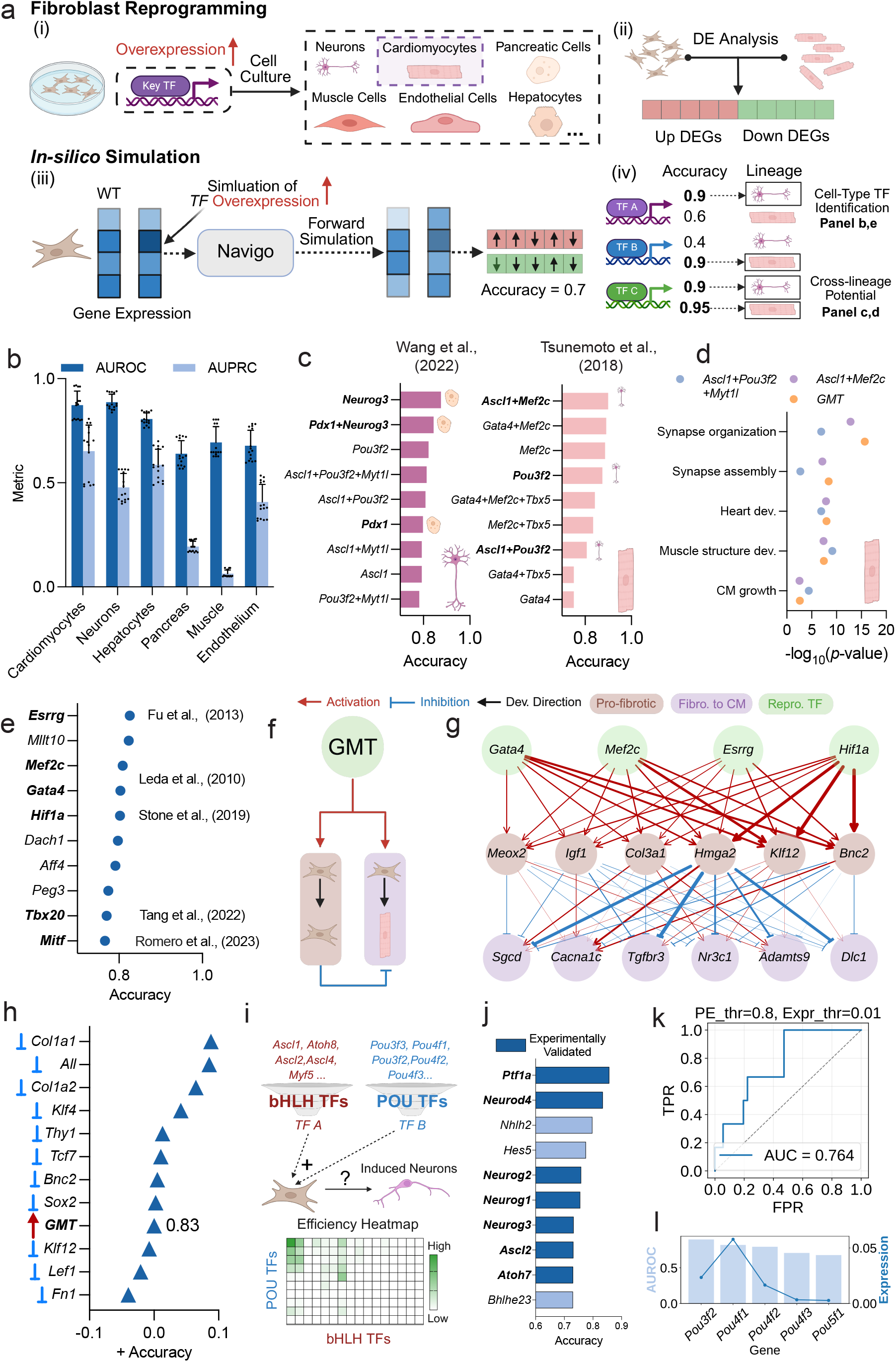
Navigo interprets and discovers key TFs for cell reprogramming. **a**. Illustration of the evaluation setup. (i) Fibroblasts can be reprogrammed into diverse cell types through overexpression of specific TFs, offering significant potential for regenerative medicine applications. (ii) Differential expression analysis identifies genes up- or down-regulated between fibroblasts and target cell types. (iii) Navigo identifies candidate reprogramming TFs via in silico simulation by modeling TF overexpression and predicting the resulting gene expression changes. (iv) Reprogramming accuracy is measured by the consistency of the direction of predicted expression changes with that of experimentally observed changes in the top 200 upregulated and 200 downregulated DEGs. This metric serves as a proxy for re-programming potential, because TFs with greater reprogramming potential are expected to induce transcriptional changes more consistent with successful target fate conversion. **b**. Quantitative evaluation of predicted reprogramming factor rankings for different target cell types. The experiments are replicated using three checkpoints with five seeds each. **c**. Navigo independently recovers experimentally validated cross-lineage reprogramming TFs from two prior studies. TFs or TF combinations previously shown to drive cross-lineage reprogramming are highlighted in bold and are ranked highly by Navigo for the corresponding target fate. **d**. Shared pathways activated by GMT (*Gata4, Mef2c, Tbx5*) TFs and established neuronal reprogramming factors. We used hypergeometric tests to obtain the p-value. Points for each pathway are jittered along the x-axis for visualization. **e**. Screening highly expressed TFs in cardiomyocytes identifies novel candidates for cardiac reprogramming from fibroblast cells. **f**. GMT TFs may simultaneously activate both reprogramming and pro-fibrotic markers initially, with pro-fibrotic signals subsequently inhibiting the reprogramming process. **g**. GRNs inferred by Navigo illustrating interactions between reprogramming TFs, pro-fibrotic markers, and cardiac development-related genes. **h**. Navigo predicted inhibition of pro-fibrotic pathway genes can lead to reprogramming efficiency improvement. **i**. Systematic screening for neuronal reprogramming TFs. Navigo’s predictions were evaluated using the experimental reprogramming efficiency map from [31]. **j**. Navigo-based ranking of bHLH TFs (bold font indicates experimentally validated successful neuronal reprogramming factors). **k**. Overall AUROC scores and performance curves for all bHLH-POU TF combinations with programming effect threshold 0.8 and expression threshold 0.01. The programming effect threshold sets measured values below a cutoff to zero, and the expression threshold selects TFs above minimum average expression. **l**. Relationship between expression level and predicted accuracy for POU TFs.

### Navigo Accurately Maps the Mouse Embryogenesis Process

A practical generative model of embryogenesis must learn the underlying regulatory process generating the observed data, rather than merely fitting static temporal snapshots. This distinction is critical because experimental scRNA-seq atlases have inherent limitations: data are collected at discrete time intervals with inevitable technical noise and sampling variability that can obscure true biological signals, particularly for cell types that are hard to capture. A model that genuinely captures the generative process should be able to infer biologically meaningful cellular states at unobserved intermediate time points (interpolation) and distinguish genuine developmental dynamics from technical artifacts (denoising). We therefore evaluated Navigo on these two tasks (Figure 2a), which assess whether the learned vector field represents a faithful reconstruction of the continuous regulatory dynamics governing embryonic development.

We first evaluated the interpolation performance of Navigo by testing whether the model accurately predicts gene expression distributions at held-out intermediate time points. We designed two prediction strategies: forward simulation, which predicts a held-out time point by integrating cells forward from an earlier observed population along the learned developmental vector field, and aligned interpolation, which predicts the intermediate population by interpolating between aligned cell pairs from the two flanking time points (*Methods*). Here we report results using aligned interpolation. Navigo consistently outperformed all baseline methods (Moscot [24], TIGON [25], and Random) across different interpolation time intervals (Figure 2b), as measured by Wasserstein distance with statistical significance confirmed by Wilcoxon matched-pairs tests. The visualization of UMAP further confirmed these results (Figure 2c), showing that Navigo-generated cells at E14.25 exhibited substantially better overlap with the ground truth compared to baseline methods. Performance for other metrics, forward simulation results, and visualizations for other time points are shown in Supplementary Figures 3 and 4.

We further evaluated how accurately models predict cell type composition during interpolation (*Methods*). Navigo demonstrated significant superiority over baseline methods across all tested time intervals, quantified with Jensen-Shannon divergence between predicted and ground truth cell type proportions (Figure 2d), with Wilcoxon matched-pairs tests confirming highly significant differences. Visualization of predicted cell type proportions at E16.25 (Figure 2e) showed that Navigo predictions closely matched the ground truth distribution across different cell types, whereas baseline methods exhibited notable deviations. Performance for other metrics, forward simulation results, and visualizations for other time points are shown in Supplementary Figures 3 and 5.

We then assessed the computational scalability of Navigo compared to other methods. As shown in Figure 2f, Navigo demonstrates superior scalability characteristics. TIGON [25], which employs a Neural-ODE framework operating in a relatively low-dimensional space with 10 features, requires substantially longer training time on large datasets. In contrast, Navigo with flow matching shows substantially better scalability while operating in a much higher-dimensional space (3,902 genes). This difference is critical because excessive dimensionality reduction inevitably results in information loss that compromises the model performance, as demonstrated by the inferior performance of TIGON in Figure 2b-d. The computational advantage of flow matching stems from its ability to directly regress the vector field through simulation-free objectives during training, whereas Neural-ODE methods require solving ordinary differential equations during both forward and backward passes.

We next evaluated the denoising capability of Navigo by testing whether the model can recover true cellular states when technical artifacts cause under-representation of specific populations. For this task, we employed forward simulation to generate denoised cell populations (*Methods*). We designed two complementary evaluation settings (Figure 2g). In the simulated setting, we randomly selected *X* cell type-time point pairs and retained only *Y* % of cells from each selected pair to simulate controlled noise scenarios across diverse developmental contexts. Specifically, we set *X* = 100 and *Y* = 20 in our experiments. In the real setting, we identified instances where certain cell types showed anomalously low proportions at particular developmental stages, likely reflecting technical capture difficulties or biological noise.

In the simulated setting, we first evaluate the agreement between the predicted denoised cell type proportions and the ground truth proportions without added noise. As shown in Figure 2h, the predicted proportions are highly correlated with the ground truth cell type proportions, with a high Pearson correlation indicating that our model can effectively recover cell type proportions that are highly consistent with the ground truth. In Figure 2i, we further evaluate the marker gene prediction capability of Navigo (*Methods*) by assessing how well the marker genes derived from the predicted cell population align with the marker genes derived from the ground truth cell population. Navigo achieves significantly higher F1 scores compared to the noised population, demonstrating its effective denoising performance. The performance comparisons under other hyperparameters for *X, Y* are shown in Supplementary Figure 6.

In the real setting, we focus on a case study of myofibroblasts, as shown in Figure 2j. The noisy cell type proportions exhibit a notable irregularity for the myofibroblast proportion at E18.25 (red box), which appears inconsistent with the smooth developmental trajectory observed at other stages. After applying Navigo denoising, this anomaly is corrected, resulting in a smoother temporal progression of myofibroblast proportions that better aligns with expected gradual developmental transitions during skeletal muscle development. To validate the denoising accuracy, we examine the expression patterns of several myofibroblast marker genes (Figure 2k). The Navigo-denoised cell population shows improved concordance with the expected expression dynamics of key markers, including *Cdkn1c* [34], *Synpo2* [35], *Myf6* [36], *Tnnc2* [37], *Myog* [38], and *Aldoa* [39], across the developmental time points from E18.0 to E18.75. Results across other marker genes are shown in Supplementary Figure 7a. Furthermore, pathway enrichment analysis (Figure 2l) reveals that upregulated genes identified from the Navigo-denoised population by differential expression analysis against myofibroblasts at other developmental time points (*Methods*) exhibit significantly stronger enrichment for relevant biological processes than those identified from the noisy population. Specifically, myofibroblast-related pathways, including skeletal muscle contraction [40], metabolic processes [41], stress response [42], and cellular movement [43], all demonstrate higher statistical significance (lower *p*-values) in the denoised population. Notably, these pathways show a consistent enrichment across adjacent time points (E18.0, E18.5, E18.75) but are not enriched in the noisy E18.25 data (Supplementary Figure 7b), confirming that Navigo effectively recovers biologically meaningful cell type populations by restoring the temporal continuity of pathway activity. Case studies on other cell types are shown in Supplementary Figure 7c-h.

### Navigo Enables Understanding of Lineage-specific and Disease-specific Gene Regulatory Networks

To evaluate whether the development vector field accurately captures lineage-specific GRNs, as shown in Figure 3a, we first extracted regulatory networks directly from the learned vector fields by computing the Jacobian matrix, which quantifies how changes in mature RNA counts *S*_*j*_ influence transcription rates *α*_*i*_ as 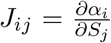. This Jacobian element *J*_*ij*_ represents the regulatory strength of Gene *j* on Gene *i*. The inferred regulatory network was then compared against ChIP-seq experimental data from the ChIP-Atlas [44] database, which measures transcription factor (TF) binding across diverse cell lineages and serves as experimental reference for validation. As shown in Figure 3b, Navigo consistently out-performed existing methods (CellOracle and Dynamo) in recovering known TF-target relationships as measured by AUROC scores (*Methods*), with statistical significance confirmed by Wilcoxon matched-pairs tests (*p*-value *<* 0.05). These results demonstrate that Navigo effectively learns the regulatory logic across different cell types.

We next applied this validated framework to investigate the gene regulatory mechanisms underlying developmental diseases by analyzing CHD [45] (Figure 3c), structural abnormalities of the heart that develop during fetal development and represent the most common type of birth defect. CHDs encompass diverse subtypes including atrial septal defects (ASDs), ventricular septal defects (VSDs), outflow tract (OT) defects, heterotaxy, functional single ventricle, and obstructive lesions. Although genes associated with different CHD subtypes are known to involve distinct developmental processes, how different CHD genes affect specific regulatory pathways and networks to cause particular defect types remains incompletely understood.

To quantify the impact of individual genes on the developmental regulatory network, we simulated knockouts of CHD-associated genes from the CHDgene database [28] by setting their expression to zero and predicting downstream transcriptomic changes using the learned vector field (*Methods*). For each knockout, we compared the perturbed trajectory to the wild-type trajectory to obtain a response vector quantifying the gene’s regulatory impact. This approach enabled systematic comparison of network perturbations caused by different CHD genes.

To obtain an overview of how different CHD genes perturb the regulatory network, we performed unsupervised clustering of the response vectors using *K*-means. We determined the optimal number of clusters using the elbow method (Supplementary Figure 8a), which identified four distinct functional modules (Figure 3d, Supplementary Figure 9). To characterize the biological processes affected by each module, we computed the average response vector for each cluster and performed pathway enrichment analysis using the hypergeometric test (*Methods*). This analysis revealed that each CHD gene cluster has distinct biological functions. Cluster 1 genes primarily affected cardiac development processes including cardiac muscle tissue development, cardiac muscle contraction, and heart morphogenesis. Cluster 2 genes primarily affected vascular development processes including blood vessel morphogenesis and circulatory system development. Cluster 3 genes regulated general embryonic development processes such as chromosome organization and Wnt signaling. Notably, Cluster 0 genes influenced neuronal development processes such as synapse organization, cell junction organization, and neuron projection guidance, consistent with clinical observations of neuro-developmental impairments in CHD patients [46, 47] and genetic evidence linking CHD and neuronal developmental programs [48]. We provided the top 20 enriched pathways for each cluster in Supplementary Figure 10. In contrast, the same analysis using Dynamo (Supplementary Figure 8b-d and Supplementary Figure 11) failed to generate biologically meaningful CHD gene clusters.

We further examined the connections between genes, disease subtypes, and the identified clusters. Figure 3e shows genes with minimal distance to each cluster centroid, with bold genes representing those with minimal summation distance across all four clusters. Notably, *Gata4, Tbx5*, and *Tbx20* from Cluster 0 are known to regulate both cardiac and neuronal development [49, 50, 51, 52, 53, 54]. *Hand1* and *Hand2* from Cluster 1 are critical for both cardiac tissue and vascular development [55, 56]. Furthermore, *Son, Arid1a*, and *Igf2* from Cluster 3 are known as critical regulators of embryogenesis [57, 58, 59, 60, 61]. We also analyzed the distribution of CHD genes across clusters for different disease subtypes (Figure 3f). Each CHD subtype appeared to be caused by a mixture of genes from different clusters, reflecting the complex genetic basis of CHD. Notably, heterotaxy showed a distinct pattern, being primarily associated with Cluster 0 (neuronal) and Cluster 1 (cardiac), with minimal representation in embryonic or vascular-specific clusters. This pattern is particularly compelling as it aligns with the known mechanistic basis of heterotaxy, which primarily results from ciliary dysfunction [62, 63]. Importantly, cilia are cellular structures that coordinate signaling networks critical for both cardiac [64, 65] and neural development [66]. To further characterize the regulatory relationships among CHD genes, we constructed a regulatory interaction network where edge width indicates the mutual regulatory impact between genes (Figure 3g, *Methods*). Genes with the strongest regulatory effects included *Gata4, Gata5, Tbx5, Myh6*, and *Myh7*, which are well-known core cardiac regulators [67, 68, 69, 70, 71]. Notably, the network revealed strong interactions between the neuronal cluster (Cluster 0) and both the cardiac cluster (Cluster 1) and vascular cluster (Cluster 2). This observation aligns with the established neurovascular congruency during development [72, 73, 74] and the crosstalk between neural and cardiac systems during development and function [75, 76]. Together, these findings validate the biological relevance of our predicted gene clusters and their associations with specific developmental processes and disease phenotypes.

We next examined pathway enrichment patterns for individual CHD types. We performed pathway enrichment analysis using the response vectors within each CHD class, calculating the ratio of genes showing significant enrichment (*p*-value *<* 0.05) and identifying pathways with the highest ratio difference between each CHD disease group and the rest of the genes. As shown in Figure 3h, CHD genes associated with OT defects showed distinctly higher enrichment in several developmental pathways compared to other CHD genes. Cell junction-related pathways showed prominent enrichment, consistent with the critical role of adherens junctions and cell-cell contacts in epithelial formation during OT development [77, 78]. Migration-related pathways, including smooth muscle cell migration and locomotion, were also enriched, reflecting the importance of cardiac neural crest cell migration and smooth muscle cell migration in OT morphogenesis [79, 80]. Additionally, differentiation and signaling pathways, including regulation of cardiomyocyte differentiation, cAMP response, and intracellular signaling, showed differential enrichment, reflecting the importance of coordinated cellular differentiation and signaling cascades in OT development [81, 82]. These pathway enrichment patterns align well with known biological mechanisms underlying OT defects, demonstrating that our model accurately captures disease-specific regulatory pathways. The results for other CHD subtypes are shown in Supplementary Figure 12a.

We then investigated whether our method can distinguish between ASD and VSD, two similar septal defects affecting different heart chambers. We calculated the impact ratio of each CHD gene on ventricular versus atrial markers (*Methods*), comparing genes causing ASD, VSD, or both defects. The differentially expressed genes (DEGs) between ventricular and atrial cardiomyocytes are shown in Supplementary Figure 12b. As shown in Figure 3i, VSD genes showed higher impact on ventricular markers while ASD genes showed higher impact on atrial markers, aligning with their anatomical locations, and genes causing both defects showed intermediate patterns. In contrast, Dynamo failed to capture this directionality. We further examined specific gene examples in Figure 3j, showing the top affected ventricular and atrial markers for representative CHD genes associated with ASD (*Smad3*), VSD (*Gata6*), and both defects (*Myh7*), which confirmed the chamber-specific regulatory effects predicted by our method. These results demonstrate that our method accurately distinguishes the cell-type-specific regulatory mechanisms underlying closely related CHD subtypes.

### Navigo Predicts Genetic Perturbation Outcomes Without Training on Specific Knockout Data

A key promise of learning developmental dynamics is the ability to perform *in silico* genetic perturbations, creating “virtual embryos” to predict how genetic knockout (KO) reshapes cell fates and gene expression programs without requiring costly and time-consuming animal work, gene editing and scRNA-seq experiments. We evaluated Navigo’s ability to predict genetic KO outcomes without training on any knockout data using an independent scRNA-seq dataset of mouse embryonic gene expression following genetic KO (Figure 4a). This dataset [29] includes six KO genotypes, generated via standard gene-editing and breeding or tetraploid aggregation, and profiled by single-cell sequencing at E13.5 alongside wild-type controls.

To assess zero-shot KO prediction, we simulated KO by replacing each KO gene’s expression in all wild-type cells with *mg*_*i*_, where *g*_*i*_ is the mean non-zero expression of gene *i* in wild-type cells, and *m <* 0 is a hyperparameter controlling the KO strength. The negative intervention value was used to represent a sustained inhibitory signal during simulation, following the perturbation strategy used in Dynamo [23]. We then applied Navigo’s ODE-based inference to predict developmental gene expression changes and identified the most affected genes by comparing the predicted KO and WT profiles (*Methods*). These predictions were validated against ground-truth DEGs, determined using a Wilcoxon rank-sum test with Benjamini-Hochberg correction (FDR *<* 0.05). Detailed statistics for different cell types and KOs are shown in the Supplementary Figure 13-15.

As demonstrated in Figure 4b, this prediction task poses several challenges. First, the two datasets originate from independent studies, creating a data gap that tests model generalization. Second, analysis of ground-truth DEGs revealed substantial variance across KO-lineage combinations, with a substantial proportion of combinations producing relatively few DEGs. Third, a temporal mismatch exists: while actual KOs occur at embryogenesis onset, data availability and computational constraints limit our simulations to nearby time points around the E13.5 profiling stage. Specifically, we used five time points (E12.5, E12.75, E13.0, E13.25, and E13.5) to simulate KO impacts and averaged the predicted gene expression changes across these time points.

Gene expression changes following KO vary across developmental lineages: some cell types exhibit substantial expression changes, while others remain relatively stable due to compensatory mechanisms. We assessed whether Navigo captures this heterogeneity by examining the correlation between predicted mean absolute expression changes and the number of ground-truth DEGs. As shown in Figure 4c and Supplementary Figure 16a, we observed a positive correlation across gene-lineage combinations (each dot represents one combination). This demonstrates that the model effectively prioritizes combinations likely to produce observable experimental outcomes, with lineage-specific correlations shown in Supplementary Figure 16c.

We next benchmarked Navigo against existing methods across different developmental lineages for *Gli2* KO. We evaluated prediction performance using Top-50 precision, defined as the percentage of ground-truth DEGs recovered among the top 50 genes ranked by predicted expression change magnitude. As shown in Figure 4d, Navigo consistently outperformed both the random baseline and CellOracle. For the random baseline, we randomly sampled a set of genes with the same size as the ground-truth DEG set for each gene-lineage combination and computed Top-50 precision against this random set. Notably, Navigo achieved higher prediction precision on gene-lineage combinations with stronger perturbation signals. Similarly, we also observed that Navigo achieved higher prediction precision in lineages with stronger perturbation signals, as measured by the number of ground-truth DEGs (Figure 4e, Supplementary Figure 16b). To assess sensitivity to the perturbation hyperparameter *m*, which controls the magnitude of simulated gene expression reduction upon KO, we evaluated Navigo’s performance across different *m* values (Figure 4f). Navigo maintained stable Top-50 precision across a range of *m* values, indicating that its prediction performance is relatively insensitive to this hyperparameter choice.

Beyond quantitative benchmarking, we investigated whether Navigo provides mechanistic insights into the variable KO responses across lineages. Genetic compensation [83, 84] represents a fundamental yet incompletely understood phenomenon in developmental biology, whereby the loss of a gene can be buffered by functional redundancy from other genes within the same gene family. As shown in Figure 4g, despite *Gli2* being an essential TF regulating multiple critical developmental pathways [85, 86, 87], *Gli2* KO embryos remain viable at E13.5, suggesting that other *Gli* family members (*Gli1* and *Gli3*) may compensate for *Gli2* loss [88, 89]. Consistent with this principle, Figure 4b shows that KO responses vary substantially across both genes and developmental lineages, indicating context-dependent compensation.

To assess whether Navigo reliably captures pathway-level compensation patterns, we performed pathway enrichment analysis on both model-predicted and experimentally observed DEGs from *Gli2* KO using the hypergeometric test. This analysis revealed heterogeneous compensation: while some developmental pathways (EMT [90], WNT [87], Hedgehog [85]) showed significant enrichment following *Gli2* KO (*p <* 0.05), others remained stable, as shown in Figure 4h (limb mesenchyme). The model-predicted enrichment patterns closely matched experimental observations, accurately identifying which pathways were significantly perturbed or stable, supporting its reliability for downstream interpretation. Results on other cell types are shown in Supplementary Figure 17.

We next analyzed the factors underlying lineage- and gene-dependent variability in KO responses. We hypothesized that, within a gene family, the magnitude of the experimental KO response depends on the perturbed gene’s relative regulatory contribution compared with its paralogs in a given lineage. To test this idea, we performed systematic KO simulations for all genes in each family and quantified, for each lineage, a family-normalized score for each member’s predicted regulatory contribution. We then compared these simulation-derived scores with DEG counts measured in the corresponding experimental KO datasets. In the *Gli* family (Figure 4i), the lineage-specific pattern of predicted *Gli2* contribution was concordant with the pattern of DEG counts across lineages in the experimental *Gli2* KO data. Specifically, lineages in which *Gli2* was predicted to contribute more strongly than *Gli1*/*Gli3* showed larger experimental KO responses (more DEGs), whereas lineages in which *Gli2* was predicted to play a weaker relative role showed smaller responses. Thus, although the simulated contribution score and experimental DEG count are different quantities, their agreement across lineages indicates that the model captures biologically meaningful differences in *Gli2* dependence. In the *Scn* family (Figure 4j), *Scn9a* consistently showed the highest predicted regulatory contribution relative to *Scn10a* and *Scn11a* across all examined lineages. Consistent with this prediction, experimental double KO of *Scn10a/11a* resulted in minimal DEGs, suggesting that *Scn9a* buffers their loss. Together, these observations support the idea that the balance of regulatory contribution among paralogs determines KO response magnitude.

Beyond explaining response magnitude, we investigated which specific downstream genes are sensitive to KO by examining regulatory directionality among family members. We hypothesized that genes co-regulated in the same direction by family members would be more sensitive to gene KO perturbations than genes subject to opposing regulation. Using Navigo, we simulated *Gli1*/*Gli3* double KO and *Gli2* KO separately and classified predicted *Gli* effectors into “Same Directions” (concordant regulation by *Gli2* versus *Gli1*/*Gli3*) and “Different Directions” (opposing regulation) categories (Figure 4k, left). Hypergeometric enrichment analysis revealed significant enrichment of experimental *Gli2* KO DEGs in the “Same Directions” category across lineages (Figure 4k, right). This pattern is consistent with dosage sharing in redundant gene regulation, where loss of one family member reduces combined regulatory output below a functional threshold, making expression changes more likely to manifest [91]. Conversely, effectors in the “Different Directions” category tend to remain buffered, as the opposing regulatory activities of remaining family members can partially compensate for the loss [92]. Supplementary Figure 18 shows similar enrichment patterns for simulated single KOs of *Gli1* and *Gli3*. Together, these results suggest that Navigo captures interpretable regulatory logic linking two complementary aspects of gene family compensation to KO outcomes: the balance of regulatory activity among paralogs may influence response magnitude, while regulatory directionality may determine which specific downstream genes are most affected.

### Navigo Interprets and Discovers Key TFs for Cell Reprogramming

Cellular reprogramming has emerged as a versatile platform for regenerative medicine, enabling direct generation of therapeutically relevant cell types such as cardiomyocytes for cardiac repair [93], neurons for neurodegenerative disease treatment [94], hepatocytes for liver regeneration [95], and insulin-producing beta cells for diabetes therapy [96]. While Navigo was trained on natural developmental trajectories from time-lapse scRNA-seq data, we hypothesized that its learned regulatory dynamics could generalize to predict induced reprogramming outcomes, as both processes involve coordinated changes in GRNs that drive cell fate transitions. Successfully predicting reprogramming outcomes would demonstrate Navigo’s practical utility for rational design of cellular reprogramming and TF discovery in regenerative medicine.

Fibroblasts can be reprogrammed into specific cell types by overexpression of key TFs in cell culture. As shown in Figure 5a, we computationally simulated this process by setting a candidate TF’s expression to a high value and performing forward inference to predict the resulting gene expression changes. We then compared the direction of these predicted changes with that of the known DEGs distinguishing the target cell type from fibroblasts, and used the resulting directional accuracy as a proxy for reprogramming potential, reasoning that TFs with stronger reprogramming potential should induce transcriptional changes more consistent with those observed during successful target fate conversion (*Methods*). First, we evaluated Navigo’s performance in predicting cell-type-specific reprogramming TFs across diverse lineages. We curated the most widely recognized TFs for reprogramming fibroblasts to specific cell types (Supplementary Figure 19a). For each target cell type, we classified all combinations of its widely recognized TFs as positive examples, while treating TF combinations for other cell types as negative examples (*Methods*). We then calculated AUROC scores based on Navigo’s prediction accuracy. As shown in Figure 5b, both AUROC and AUPRC scores were consistently high across all lineages, demonstrating robust and accurate predictive capabilities of our model. To ensure result reliability, we validated these findings using three different model checkpoints for each cell type and bootstrapped five different subsets for each cell type, further confirming the robustness of our predictions.

Interestingly, some high-ranking TFs in Navigo are not canonical regulators of the target lineage. Rather than reflecting model limitations, these predictions may capture genuine cross-lineage reprogramming potential [30]. While early reprogramming studies [97, 98] assumed TFs possessed unique lineage-specific capabilities, recent work [30] has revealed that certain TFs function as cross-lineage regulators capable of driving the reprogramming of multiple distinct cell fates. To investigate whether Navigo could independently discover such factors through *in silico* simulation, we analyzed predicted accuracy rankings for cardiomyocytic and neuronal reprogramming (Figure 5c). Notably, the neuronal factor *Ascl1* ranked highly in cardiomyocytic predictions, surpassing canonical GMT factors (*Gata4, Mef2c, Tbx5*), consistent with experimental evidence of *Ascl1*’s cross-lineage functionality [30]. Similarly, Navigo independently identified the neuronal reprogramming potential of *Neurog3*, initially characterized for fibroblast-to-pancreatic beta cell conversion [99] and later validated for neuronal reprogramming [31]. To further validate Navigo’s predictions, we compared perturbation predictions of top-ranked TF combinations against experimental results [30]. Pathway enrichment analysis (Figure 5d) of highly activated downstream genes showed strong concordance: both GMT and neuronal reprogramming TFs activated overlapping pathways in synapse organization, heart development, and muscle structure development, demonstrating the biological relevance of the cross-lineage regulatory mechanisms predicted by our approach.

We then evaluated Navigo’s capabilities in systematically screening a wider range of TFs to identify alternative candidates for cell reprogramming. Cardiomyocytes were selected as the target cell type due to extensive research in this field. Initially, we selected the top 50 TFs with the highest expression levels in cardiomyocytes, followed by screening and ranking these TFs based on prediction accuracy. As shown in Figure 5e, we identified the top 10 most promising TFs for reprogramming, with TFs validated by existing literature highlighted in bold text. For instance, *Esrrg*, when combined with GMT factors and applied to human fibroblasts, significantly enhances reprogramming success [100]. Similarly, *Tbx20* promotes cardiac reprogramming by activating genes associated with cardiac contractility, maturation, and ventricular development [101]. Other literature-validated factors include *Hif1a* [102] and *Mitf* [103, 104]. These findings demonstrate Navigo’s potential for identifying promising TF candidates even without prior knowledge of GRNs.

Programming efficiency represents a primary challenge in fibroblast-to-cardiomyocyte reprogramming for practical applications [105]. Recent studies have identified that GMT factors concomitantly activate pro-fibrotic pathways during cardiac reprogramming, which subsequently inhibit fibroblast conversion [105] (Figure 5f). We sought to recapitulate this regulatory mechanism using Navigo. By systematically simulating overexpression of candidate genes and quantifying their regulatory impacts, we reconstructed regulatory relationships among key genes involved in this process (Figure 5g). Our analysis revealed a regulatory cascade where cardiac-promoting TFs (*Gata4, Mef2c, Esrrg, Hif1a*, green nodes) activate pro-fibrotic markers (*Meox2, Igf1, Col3a1, Hmga2, Klf12, Bnc2*, brown nodes), which in turn suppress essential cardiac developmental genes (*Sgcd, Cacna1c, Tgfbr3, Nr3c1, Adamts9, Dlc1*, purple nodes). This Navigo-inferred network recapitulates the experimentally observed antagonistic mechanism. Furthermore, *in silico* knockdown of the pro-fibrotic genes along with the overexpression of GMT genes substantially improved Navigo’s prediction accuracy (Figure 5h), consistent with experimental findings that pro-fibrotic pathway suppression enhances reprogramming efficiency [105], thereby supporting the biological relevance of our computational model.

We next evaluated Navigo’s capacity to predict fine-grained reprogramming outcomes by examining combinations of established neuronal reprogramming factors from two well-characterized TF families: bHLH TFs [106] and POU TFs [107]. We leveraged data from a recent study [31] that systematically screened pairwise combinations across 46 bHLH family members and 12 POU family members, generating quantitative reprogramming efficiency maps that classified TF pairs as either successful neuronal inducers or failed combinations (Supplementary Figure 19b). Following the same combinatorial screening strategy employed experimentally, we computationally evaluated all possible TF pairs using Navigo to predict their reprogramming efficiency (Figure 5i). This framework presents a stringent test of Navigo’s predictive capabilities due to two challenges: the relatively small functional distinctions within the same TF family and the need to predict synergistic interactions between factor pairs rather than individual effects.

We first conducted systematic screening of bHLH TFs, where only 16 of the 46 family members demonstrated successful experimental reprogramming capabilities. Single TF overexpression screening identified the top 10 candidates most likely to facilitate fibroblast-to-neuron reprogramming as predicted by Navigo (Figure 5j). Remarkably, 7 of these 10 top-ranked TFs were experimentally validated (bold) to induce neuronal programming, demonstrating our model’s capability to effectively prioritize promising candidates within the same TF family.

For combinatorial screening, we applied two filtering thresholds to improve prediction robustness. The experimental study [31] quantified neuronal induction for each TF pair as the percentage of TUJ1-positive cells among plated fibroblasts at day 14 post-induction, with higher values indicating stronger reprogramming activity. We therefore applied a programming effect threshold to this experimental read-out, setting values below a cutoff to zero to exclude weak or potentially noisy effects. In parallel, the expression threshold filtered lowly expressed TFs that may be more challenging to predict accurately. As shown in Figure 5k, Navigo achieved an AUROC of 0.764. We further evaluated different combinations of programming effect and expression thresholds (Supplementary Figure 20) and found that the model maintained reasonable performance across most parameter settings, with improved performance under stricter thresholds. We also evaluated anchor-based combinatorial screening, where one TF from a given family was fixed while screening the other family for optimal partners, and observed similarly robust performance across a broad range of threshold settings, with generally improved AUROCs under more stringent filtering (Supplementary Figure 21).

We next examined how model performance varies with TF expression levels (Figure 5l and Supplementary Figure 22). By calculating the mean expression level of each TF and its corresponding AUROC in anchor-based combinatorial screening, we observed a positive correlation, indicating better predictions for highly expressed TFs. This suggests that predicting reprogramming outcomes for lowly expressed TFs remains challenging in zero-shot settings, where limited expression signal may tend to reflect technical noise and thus hinder accurate modeling. Improving performance on lowly expressed TFs represents a promising direction for future work.

## Discussion

In this paper, we introduce Navigo, a biologically grounded generative modeling framework that learns predictive developmental dynamics from static snapshots of embryogenesis. This framework addresses a fundamental challenge in developmental biology: while single-cell technologies have generated unprecedented atlases profiling millions of cells across development [13, 14, 15, 16], these data capture discrete snapshots rather than the continuous dynamical system that generates cell-fate transitions. Understanding these dynamic regulatory processes is essential for building generalizable AI platforms that can model developmental diseases, predict the consequences of genetic and regulatory perturbations, and guide rational cell-fate engineering for regenerative medicine.

Navigo advances this vision by integrating flow matching at the population level with RNA kinetics modeling at the molecular level, learning continuous dynamics that govern cell state transitions during development. We evaluate Navigo along three complementary axes that reflect its potential as a generalizable AI platform for developmental biology. First, in disease modeling, Navigo reveals functionally distinct gene modules underlying different CHD subtypes, identifying pathway signatures and chamber-specific regulatory effects consistent with known disease biology. Second, in perturbation-effect prediction, Navigo achieves accurate zero-shot prediction of six knockout phenotypes on an independent dataset, uncovering lineage-specific compensation mechanisms in which predicted relative regulatory strength within gene families correlates with experimental knockout severity. Third, in rational cell-fate engineering, Navigo successfully identifies established reprogramming factors, reveals pro-fibrotic barriers to cardiac reprogramming, discovers cross-lineage regulators, and achieves high accuracy in predicting hundreds of transcription factor combinations effective for neuronal conversion. Together, these results position Navigo as a generalizable AI platform that transforms static developmental atlases into predictive systems for disease modeling, perturbation-effect prediction, and rational cell-fate engineering.

Despite these advances, several limitations suggest important directions for future improvement. First, while our use of metacells substantially reduces computational cost compared with modeling individual cells, and flow matching is more efficient than neural ODEs [108, 109], Navigo still requires substantial training time. Developing more scalable model architectures and optimization strategies would improve efficiency and enable broader adoption of generative modeling approaches in developmental biology. Second, our perturbation prediction is based on zero-shot inference and validated on observational knockout data without direct supervision from perturbation experiments. Although this demonstrates the model’s ability to learn meaningful regulatory relationships, fine-tuning or explicitly training on perturbation datasets could further improve prediction accuracy, particularly for subtle or context-dependent perturbation effects. Third, due to data availability, our analysis focuses on development from E8.5 onward [13], missing earlier critical stages of embryogenesis, including pre-implantation development and gastrulation. Training models on datasets spanning the full developmental timeline would enable a more holistic understanding of embryogenesis. Ideally, such datasets should be generated using unified experimental platforms to minimize batch effects and ensure comparability across stages. Fourth, although Navigo models developmental state transitions in transcriptional space and succeeds in many applications, it does not explicitly account for population dynamics such as cell proliferation, division, and death, nor does it incorporate lineage information or ancestor–descendant relationships. These processes are fundamental to embryonic development and may shape both the directionality and abundance of emerging cell states. Integrating lineage tracing and explicit population dynamics into the modeling framework could improve the biological realism of the learned developmental trajectories and strengthen mechanistic interpretation.

Several promising directions emerge for advancing predictive and programmable modeling of developmental processes. First, scaling generative models across multiple species including human, mouse, and other model organisms could reveal conserved regulatory principles and evolutionary discrepancies in developmental programs [110, 111]. Comparative analysis across species would uncover universal gene regulatory mechanisms that are essential for development while identifying lineage-specific adaptations that generate phenotypic diversity. Second, establishing active learning frameworks [112] that create feedback loops between computational predictions and genetic perturbation experiments could iteratively refine perturbation prediction accuracy while systematically validating and discovering novel regulatory mechanisms during development. Third, incorporating spatial information into the modeling framework from technologies like spatial transcriptomics [113, 114, 115, 116] would capture the critical interplay between gene regulation and spatial organization during development, enabling models to predict not only cellular identities but also tissue architecture and morphogenesis. Fourth, integrating multi-modal and multi-omics measurements [117, 118, 119] such as chromatin accessibility and DNA methylation would provide a more complete mechanistic understanding of how regulatory programs orchestrate cell fate specification during embryo development.

Looking forward, these advances point toward an ambitious long-term vision: the construction of comprehensive virtual embryo models [12, 120] that simulate complete developmental processes *in silico*. Such virtual embryos would enable simulation of entire developmental trajectories from fertilization to organogenesis, integrating gene regulation [121], cellular dynamics [122], and tissue morphogenesis [123]. Therefore, Navigo has the potential to shift developmental biology from retrospective observation to prospective prediction, enabling researchers to computationally simulate and manipulate developmental processes and fundamentally reshape therapeutic discovery for developmental disorders.

## Methods

We adopt the following notation throughout the Methods Section. Let *T* denote the number of discrete time points during embryogenesis, with actual time values *t*_1_, *t*_2_, …, *t*_*T*_ . For each time point *t*_*i*_:

- **X**_*i*_ denotes the complete cell population at time *t*_*i*_, *N*_*i*_ = |**X**_*i*_| is the number of cells and *M* is the number of genes, with each row representing the gene expression profile of a cell.
- 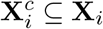 denotes the subset of cells belonging to cell type *c* at time *t*_*i*_.
- 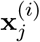 denotes the *j*-th cell in population **X**_*i*_, representing its gene expression profile at time point *t*_*i*_.
- **x**_*j*_(*t*) denotes the gene expression state of cell *j* at continuous time *t*, obtained by integrating the velocity field from an initial condition. At discrete observation times, we assume the integrated expression matches with the observed data, e.g., 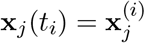.
- For model training and inference, each cell state is represented by its spliced and unspliced mRNA counts: 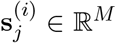 and 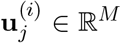. The concatenated representation 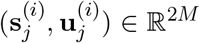 serves as input to the velocity field network.

Given temporal cell populations 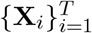, we aim to construct a model **v**(**x**, *t*) that captures developmental dynamics, described by:

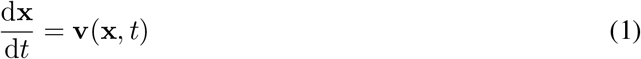

where **v** is parameterized by a neural network and represents the vector field of embryonic development. For each time point *t*_*i*_, we define the predicted distribution 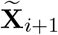 by evolving cells from **X**_*i*_:

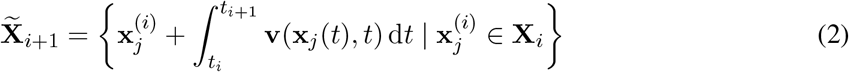

We optimize the vector field by minimizing the distributional distance between observed and predicted distributions:

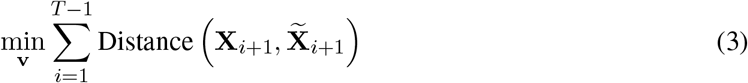

### The Navigo Framework

#### Model Architecture

The vector field network **v**(**x**, *t*) takes the cellular state information, specifically the smoothed unspliced (**u**) and smoothed spliced (**s**) gene counts, through multiple multilayer perceptron (MLP) layers to predict the fundamental kinetic parameters of RNA dynamics, ***α, β***, and ***γ***. These parameters define the velocity field through the RNA velocity equations [22].

Specifically, the transcription rate ***α*** = ***α***(**s**, *t*) is modeled as a function of spliced mRNA counts, capturing the principle that gene transcription is regulated by the expression levels of upstream regulators. Therefore, each element *α*_*i*_ is both cell-specific and gene-specific, reflecting the heterogeneous regulatory states across cells. In contrast, the splicing rates ***β*** and degradation rates ***γ*** are gene-specific constant vectors, as they are primarily determined by the intrinsic biochemical properties of each gene [32].

The vector field output comprises two components, **v**_*u*_ and **v**_*s*_, computed as:

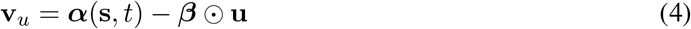

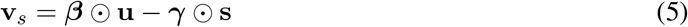

where ⊙ denotes element-wise multiplication.

This formulation is biologically grounded and designed to capture the underlying molecular dynamics of gene expression, where transcriptional regulation drives cellular state changes during development.

#### Model Training

While the objective of minimizing distributional distance between observed and predicted cell populations is theoretically sound, it encounters significant computational challenges when applied to embryogenesis data. A natural approach would be to predict future cell states by numerically integrating the 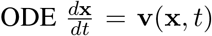 for each cell from time *t*_*i*_ to *t*_*i*+1_, compute the distributional distance between predicted and observed populations, and backpropagate gradients through the ODE solver using the Neural Ordinary Differential Equation (Neural ODE) framework [124]. However, this becomes computationally intractable for embryogenesis data characterized by high dimensionality (thousands of genes), multiple time points, and large sample sizes (hundreds of thousands of cells), as gradient computation requires solving an adjoint ODE backward in time for each training iteration.

Inspired by the flow matching method [26], we develop a computationally efficient approach for learning developmental dynamics from multi-timepoint scRNA-seq data. The core idea is an iterative refinement procedure that alternates between constructing cell pseudo-pairs and training the vector field to match velocities along linear interpolation paths connecting these pairs.

We initialize the procedure with random pseudo-pairs [26]. For each cell 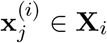 at time point *t*_*i*_, we randomly select a paired cell from the subsequent time point:

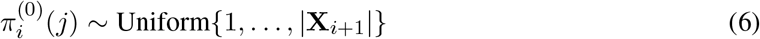

where 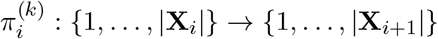 denotes the pairing function at iteration *k* that maps cell index *j* in **X**_*i*_ to its pseudo-paired cell index in **X**_*i*+1_.

For each refinement round *k* = 1, 2, …, *K*, we train the vector field network using the previous pseudo-pairs 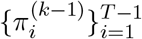 by minimizing the prediction error across all consecutive time point pairs.

For a pseudo-pair 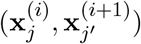 where 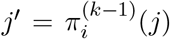, we define the interpolation path 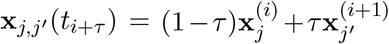 for *τ* ∈ [0, 1] and time *t*_*i*+*τ*_ = (1 − *τ*)*t*_*i*_ + *τt*_*i*+1_ . Given the cell state representation **x** = (**s, u**), the optimization objective at iteration *k* is:

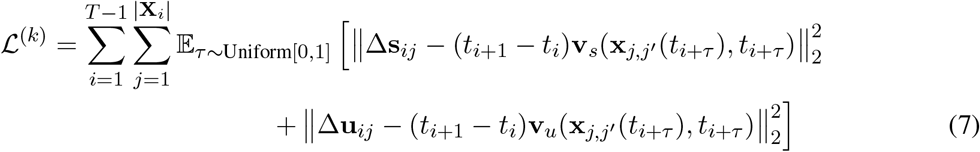

where 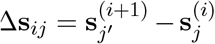 and 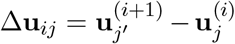 denote the displacements in spliced and unspliced gene counts, respectively.

Second, we use the trained vector field **v**^(*k*)^ to refine the pseudo-pairs. We simulate forward from each time point **X**_*i*_ to generate predicted states at time point *i* + 1. For each cell 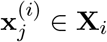, we compute:

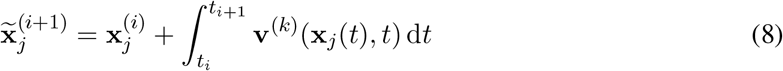

where the integral is approximated using Euler’s method with *N* = 100 discretization steps. We then update the pairing by finding the nearest neighbor in **X**_*i*+1_ for each predicted state:

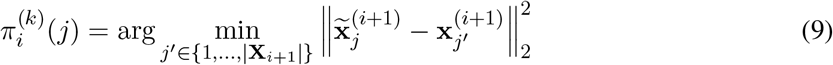

This iterative procedure progressively refines both the vector field and the pseudo-pairs, and converges to a self-consistent solution where the learned dynamics accurately predict the cell state transitions across time points [26].

#### Model Inference

##### Temporal Prediction

Navigo enables prediction of cell states at specific time points through forward integration of the learned velocity field. To predict cell states at a target time *t*_*i*+*m*_, two prediction strategies can be employed:

###### Forward simulation

Starting from an earlier observed time point *t*_*i*_, we predict cell states at target time *t*_*i*+*m*_ by integrating the learned velocity field:

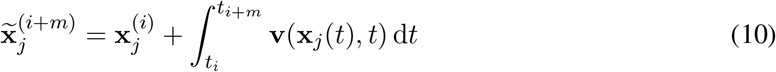

where **x**_*j*_ ∈ **X**_*i*_ and the integral is approximated using Euler’s method with *N* = 100 discretization steps. The predicted population is 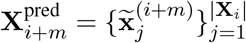. This approach requires only a single observed time point and can predict states at any future time within the model’s trained temporal range.

###### Aligned interpolation

When observations are available at both flanking time points *t*_*i*_ and *t*_*i*+*k*_, we can construct predictions by interpolating between aligned cell pairs. We first establish cell-to-cell correspondence by simulating trajectories from **X**_*i*_:

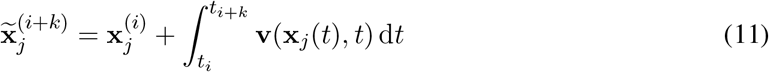

and identifying nearest neighbor alignments:

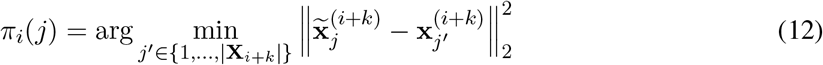

where 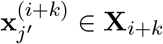 . The prediction is then obtained by linearly interpolating between aligned pairs:

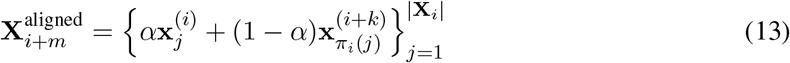

where the interpolation weight 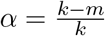 determines the relative contribution from each time point.

##### Perturbation Simulation

Navigo can simulate the effects of genetic perturbations by modifying specific gene expressions and forward integrating the learned velocity field while maintaining the perturbation. Given a cell state 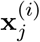 at time point *t*_*i*_, we create a perturbed state 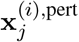 by modifying specific gene expressions while keeping other genes unchanged.

For gene *g* ∈ *G* where *G* denotes the set of genes to be perturbed, we modify the gene expression using:

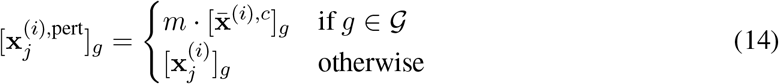

where 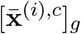 is the mean non-zero expression of gene *g* across all cells of type *c* at time point *t*_*i*_, and *m* is a perturbation strength parameter. For gene knockout, *m* ≤ 0, and for gene overexpression, *m >* 1. Multiple genes can be perturbed simultaneously by including them in *G* and applying corresponding perturbation strengths.

We then simulate both the perturbed trajectory and the wild-type trajectory by integrating the learned velocity field from time point *t*_*i*_ forward. For any time *t* ≥ *t*_*i*_:

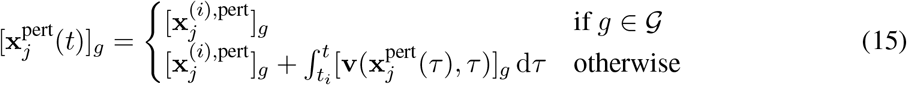

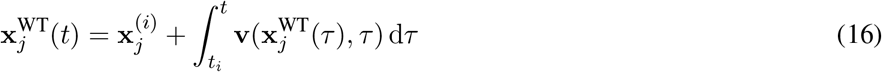

where genes in *G* remain fixed at their initial perturbed values throughout the integration, while non-perturbed genes evolve according to the learned dynamics. This approach simulates experimental scenarios such as CRISPR knockout or stable transgenic overexpression, where the intervention persists throughout the developmental timecourse.

##### Jacobian Matrix Modeling

To infer gene regulatory interactions from the learned vector field model, we compute the Jacobian matrix **J**, where each element *J*_*ij*_ represents the partial derivative of the transcription rate of effector gene *g*_*i*_ with respect to the expression level of regulator gene *g*_*j*_:

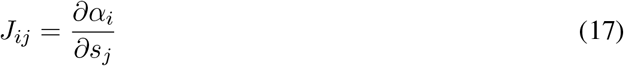

where *α*_*i*_ denotes the transcription rate and *s*_*j*_ denotes the spliced (mature) RNA counts. This Jacobian matrix quantifies the instantaneous regulatory influence of regulator expression on effector gene transcription rates as learned by the developmental vector field.

### Datasets

#### Data Collection

We used the comprehensive mouse prenatal development atlas [13], containing single-cell transcriptional profiles of 12.4 million nuclei from 83 precisely staged embryos spanning late gastrulation (E8.5) to birth (P0). This dataset offers temporal resolution at 6 hour intervals and includes annotations of over 190 cell types across major developmental lineages.

#### Data Preprocessing

We performed total count normalization and log transformation to pre-process the dataset. For downstream analysis, we retained all mouse TF genes and selected highly variable genes, retaining 3,902 genes in total. We then used scVelo [22] to compute normalized spliced (Ms) and unspliced (Mu) gene counts for RNA velocity estimation, where Ms and Mu represent locally smoothed expression levels of mature and precursor mRNA, respectively.

The transcriptomic atlas spanning the entire embryogenesis process contains tens of millions of cells. While this comprehensive dataset provides rich characterization of developmental dynamics, directly training deep learning models on such large-scale data is computationally prohibitive. To address this challenge, we adopt a metacell strategy that aggregates similar cells to reduce data size while preserving biological heterogeneity. Metacells have been widely applied in single-cell analysis tasks including trajectory inference and differential expression analysis [125, 126, 127].

We construct metacells separately for each combination of embryo, cell type, and time point. For any given embryo at a specific time point, if the number of cells belonging to a particular cell type 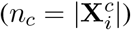 is smaller than a threshold *N* (set to *N* = 100), these cells are retained without aggregation to preserve information from rare cell populations. Otherwise, we perform *K*-means clustering with 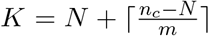, where the hyperparameter *m* is set to 50. This adaptive clustering strategy ensures that rare cell types are preserved while abundant populations are appropriately downsampled. Using this approach, the dataset size is reduced by approximately 20-fold, achieving substantially improved computational efficiency while maintaining representation of rare cell populations.

### Baselines

#### Dynamo

Dynamo [23] reconstructs continuous velocity vector fields by modeling RNA transcription, splicing, and degradation kinetics. The method estimates RNA velocities from spliced and unspliced counts and performs vector field reconstruction in reduced-dimensional space. The reconstructed vector field enables differential geometry analysis including Jacobian, acceleration, and curvature calculations for inferring gene regulatory relationships and cellular dynamics.

#### Moscot

Moscot [24] is a scalable optimal transport framework that maps cells through time and space using multi-omics single-cell data. For temporal analysis (moscot.time), the method estimates cellular trajectories by solving Wasserstein-type optimal transport problems between consecutive time points. The framework supports multimodal data through shared latent representations and achieves linear time and memory complexity through low-rank approximations of the coupling matrix, enabling application to atlas-scale datasets.

#### TIGON

TIGON [25] simultaneously infers cell velocity and population growth by solving a dynamic unbalanced optimal transport problem based on Wasserstein-Fisher-Rao distance. The method models cellular dynamics through a continuity equation with a growth term, where the velocity field describes instantaneous changes in gene expression and the growth term captures population changes due to cell division and death. TIGON uses neural networks to parameterize both velocity and growth fields, and learns trajectories by minimizing the combined Wasserstein and Fisher-Rao metrics through neural ordinary differential equations. This formulation provides an unbiased approach to learning cell transitions.

#### CellOracle

CellOracle [128] integrates GRN modeling with *in silico* perturbation to predict changes in cell identity following TF perturbations. The method constructs cell-type-specific GRN configurations using regularized linear regression models on multi-omics data, combining scRNA-seq with scATAC-seq to identify accessible promoter and enhancer regions. CellOracle simulates TF perturbations by propagating expression shifts through the inferred GRN via iterative signal propagation, then converts the simulated gene expression changes into transition probability vectors in low-dimensional embedding space.

#### Naive Baselines

##### Random

The baseline performs random cell pairing between adjacent time points and applies linear interpolation. For predicting **X**_*i*+*m*_, we randomly pair cells from **X**_*i*_ with cells from **X**_*i*+*k*_. Denoting the random pairing function as *π*_rand_, the prediction is constructed as:

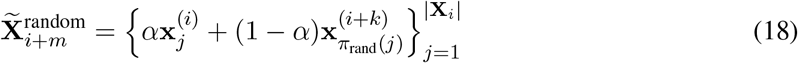

where 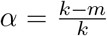 and 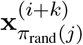 denotes the randomly paired cell from **X**_*i*+*k*_. This baseline tests whether simple temporal interpolation without learning the regulatory dynamics achieves reasonable predictions.

##### Previous

The baseline uses the cell population from the previous time point as the prediction for the target time point. Specifically, for predicting **X**_*i*_, we directly use **X**_*i*−1_ without any transformation, assuming cellular states remain unchanged over the interval.

##### Next

The baseline uses the cell population from the subsequent time point as the prediction. For predicting **X**_*i*_, we directly use **X**_*i*+1_, assuming the cells have already transitioned to their future states.

### Evaluation of Trajectory Mapping

#### Experimental Setup

We evaluate the trajectory mapping capabilities of Navigo through two complementary tasks that assess whether the learned vector field accurately captures developmental dynamics.

For temporal interpolation, we assess the ability of the model to predict gene expression distributions at unobserved intermediate time points. The complete dataset contains observations at intervals of Δ*t*_true_ = 0.25 days. We construct three experimental settings with sampling intervals of 0.5, 1.0, and 2.0 days, training the model on subsampled time points and evaluating predictions on all held-out intermediate time points. We evaluate both forward simulation and aligned interpolation performance.

For data denoising, we evaluate the ability of the model to recover true cellular state distributions when certain cell populations are underrepresented due to technical capture difficulties or sampling biases. We randomly select *X* cell type-time point pairs and retain only *Y* % of cells from each selected pair, maintaining original cell counts for all other pairs. We train the model on this biased dataset and evaluate whether forward simulation from adjacent time points can reconstruct the complete population distributions at the undersampled time points.

#### Metrics and Analysis

##### Distribution Agreement Metrics

We employ three complementary metrics to quantify distributional alignment between predicted and true cell populations at held-out time points. Lower values indicate better alignment.

###### Wasserstein Distance

The Wasserstein distance measures the minimum cost of transforming one distribution into another. For each gene *g*, we compute the 1-Wasserstein distance as

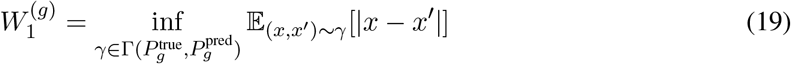

where 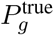 and 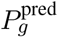 denote the empirical distributions of gene *g* in the true and predicted populations, and 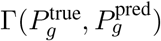 is the set of all joint distributions with marginals 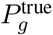 and 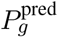. We report the average across all genes as 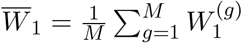.

###### Maximum Mean Discrepancy (MMD)

MMD measures the distance between distributions in a reproducing kernel Hilbert space. Using the RBF kernel 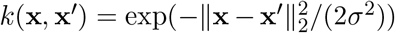, the squared MMD is computed as

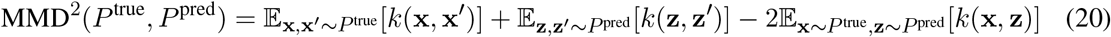

where *P* ^true^ and *P* ^pred^ represent the true and predicted population distributions. We use the default bandwidth parameter *σ* from scikit-learn.

###### Energy Distance

The energy distance is a metric between probability distributions based on expected distance between random samples. For each gene *g*, we compute

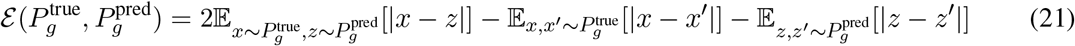

where **x, x**^′^ are independent samples from the true distribution and **z, z**^′^ from the predicted distribution. To ensure computational efficiency, we randomly subsample 1000 cells from each population when computing the energy distance for each gene. We report the average energy distance across all genes as 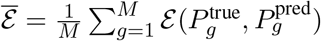.

##### Cell Type Composition Metrics

We employ two metrics that compare the distribution of cell types between true and predicted populations.

We first train a k-nearest neighbors (KNN) classifier on all training time points to predict cell type labels. Let *D*_train_ = {(**x**_*j*_, *c*_*j*_)} denote the set of cells from all training time points with their corresponding cell type labels *c*_*j*_ ∈ *C*, where *C* is the set of all cell types. We train a KNN classifier *f*_KNN_ with *n*_neighbors_ = 20 for cell type prediction on *D*_train_. We then apply this classifier to both the true population and the predicted population to obtain cell type predictions for each cell.

Let *N* ^true^(*c*) denote the number of cells in the true population predicted as cell type *c*, and *N* ^pred^(*c*) the corresponding count in the predicted population. We compute the cell type composition as probability distributions

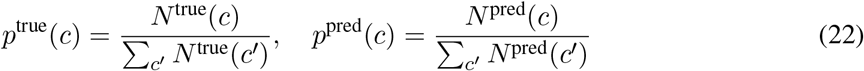

We measure the divergence between these distributions using two metrics.

###### Jensen-Shannon Divergence

The Jensen-Shannon divergence is a symmetric and bounded measure of similarity between probability distributions, defined as

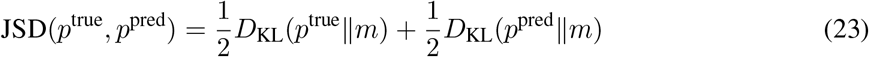

where 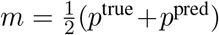 is the average distribution and *D*_KL_ denotes the Kullback-Leibler divergence. The JSD ranges from 0 (identical distributions) to 1 (maximally different distributions).

###### L1 Distance

The L1 distance, also known as total variation distance, measures the difference between the two distributions as

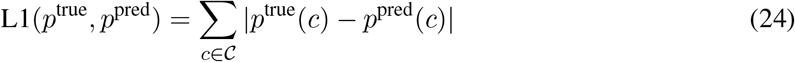

This metric ranges from 0 (identical distributions) to 2 (completely disjoint distributions), with lower values indicating better preservation of cell type composition.

##### Analysis for Cell Type Proportion in Simulated Denoising Setting

For the denoising evaluation, we apply the KNN-based cell type prediction approach to the predicted denoised population 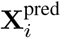. Using the trained classifier *f*_KNN_, we predict cell type labels for cells in 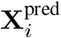 and compute the predicted cell type composition distribution *p*^pred^(*c*). For the observed biased population 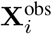 and the true complete population 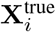, we directly use their known cell type annotations to compute the empirical distributions *p*^obs^(*c*) and *p*^true^(*c*).

##### Analysis for Marker Gene Prediction in Simulated Denoising Setting

To evaluate whether denoised predictions recover biologically meaningful gene expression signatures, we assess the ability of the model to identify DEGs that serve as cell type markers in underrepresented populations. For each cell type *c* at time point *t*_*i*_ in the complete dataset, we identify ground truth marker genes by comparing cells of type *c* at *t*_*i*_ against the same type at all other time points using the Wilcoxon rank-sum test (Benjamini-Hochberg adjusted *p <* 0.05, |logFC| *>* 0.5), yielding upregulated markers 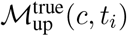 and downregulated markers 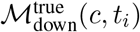. For the denoised prediction 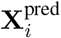, we use the trained KNN classifier to identify cells predicted as cell type *c*, denoted as 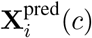, and perform differential expression analysis comparing 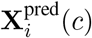 against cells of the same type from all other time points in the training data **X** _¬*i*_(*c*) using the same thresholds, yielding predicted marker sets 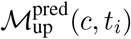 and 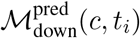.

For each direction, we compute precision, recall, and F1 score: Precision = |ℳ^pred^ ∩ ℳ^true^ |*/*|ℳ^pred^|, Recall = |ℳ^pred^ ∩ ℳ^true^ |*/*|ℳ^true^|, and F1 = 2 Precision Recall*/*(Precision + Recall). We report the average of upregulated and downregulated metrics for each cell type-time point pair.

##### Analysis for Marker Gene Signatures in Real Setting

To evaluate whether our predictions capture biologically meaningful developmental programs, we perform pathway enrichment analysis using Gene Ontology Biological Process (GOBP) terms from the Molecular Signatures Database (MSigDB v2025.1) [129].

For each predicted or ground truth cell population at a given time point, we identify DEGs by comparing the target population against all other time points of the same cell type using the Wilcoxon rank-sum test (Benjamini-Hochberg adjusted *p* < 0.01, logFC > 1). Upregulated genes are then tested for pathway enrichment using Fisher’s exact test with a one-sided alternative hypothesis.

### Evaluation of the Gene Regulatory Network

#### Experimental Setup

For evaluation of lineage-specific gene regulatory networks, we validate the inferred regulatory relationships against experimentally validated TF-target gene interactions from ChIP-seq experiments. The Jacobian matrix captures direct, first-order regulatory effects that are conceptually aligned with physical TF binding measured by ChIP-seq. To obtain stable lineage-specific estimates, we compute the Jacobian matrix at all cell states throughout the developmental trajectory of each cell type and average across all corresponding cells and time points. For each regulator, we rank all genes by the absolute magnitude of the averaged Jacobian to identify predicted regulatory targets.

We obtained experimentally validated, cell type-specific regulatory relationships from the ChIP-Atlas database [44], a comprehensive repository of processed ChIP-seq experiments, across multiple lineages including macrophages, pre-B cells, B cells, natural killer cells, cerebellum, CD4+ T cells, mast cells, heart tissue, hepatocytes, and dendritic cells. To ensure high-confidence interactions, we retained only TF-target pairs with ChIP-Atlas scores exceeding 50.

For evaluation of disease-specific regulatory mechanisms, we analyze the regulatory impact of CHD-associated genes through gene perturbation simulations. We obtained CHD-associated genes from the CHDgene database [28], a manually curated resource that catalogs genes reproducibly shown to cause CHD when mutated in humans. CHDgene employs stringent inclusion criteria, requiring that variants in each gene have been reported as the monogenic cause of CHD in at least three independent familial or sporadic cases. We focus on first heart field (FHF) and second heart field (SHF) progenitor populations, as these cardiac lineages are critical for understanding chamber-specific developmental defects.

For each CHD gene *g*, we simulate knockouts at each developmental time point *t*_*i*_ where the target cell type (FHF or SHF) is present. We set the perturbation hyperparameter *m* = 0 and predict both perturbed and wild-type cell populations at time point *t*_*i*+1_ using the perturbation simulation framework.

We define the response vector **r** by averaging the transcriptomic differences between perturbed and wild-type predictions across all cells and all time points where the target cell type exists, then normalizing to unit L2 norm:

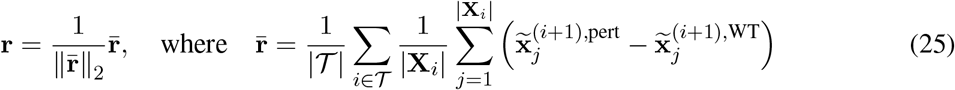

where *T* denotes the set of time point indices where the cell type of interest (FHF or SHF) is present, and ∥· ∥_2_ denotes the L2 norm. Each component *r*_*j*_ represents the normalized average change in expression of gene *g*_*j*_ caused by knockout of gene *g*. We then analyze the similarity between response vectors to identify functional modules of CHD genes with shared regulatory impacts.

To ensure lineage-specific regulatory mechanisms are properly captured, we train separate models for each major developmental trajectory. For CHD-associated gene analysis, we restrict the training set to cardiac-relevant cell populations including FHF, SHF, atrial cardiomyocytes, and ventricular cardiomyocytes.

#### Metrics and Analysis

##### Analysis of Lineage-specific GRN

For each TF, we rank all genes by the absolute value of the averaged Jacobian entries corresponding to its regulatory effects. We compute AUROC scores using this ranking, treating experimentally validated ChIP-seq targets from ChIP-Atlas as positive labels and all other genes as negative labels. Values closer to 1 indicate better discrimination of true targets from non-targets. We report the mean AUROC across all evaluated TFs for each lineage.

##### Clustering of the Response Vectors

To identify groups of CHD-associated genes with similar regulatory impacts, we perform clustering analysis on the response vectors **r**^(*g*)^ obtained from knockout simulations. We apply PCA to reduce dimensionality to 50 principal components, then apply *k*-means clustering to group CHD genes with similar downstream regulatory effects. We determine the optimal number of clusters using the elbow method, computing within-cluster sum of squares (inertia) for *k* values from 2 to 10 and selecting the *k* where the rate of decrease in inertia begins to diminish.

##### Quantifying the Mutual Regulatory Impact between CHD Genes

To investigate regulatory interactions among CHD-associated genes, we quantified the mutual regulatory impact between gene pairs. For each CHD gene pair (*g*_*a*_, *g*_*b*_), we compute their mutual regulatory strength by summing the bidirectional effects from their knockout response vectors 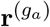 and 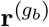:

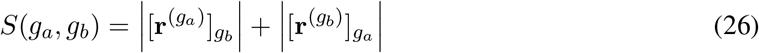

where 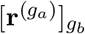 denotes the expression change of gene *g*_*b*_ in the response vector from knocking out gene *g*_*a*_. Higher values indicate stronger mutual regulatory coupling between gene pairs.

To identify the most functionally interconnected CHD genes, we rank all gene pairs by their mutual regulatory strength and retain the top 50 pairs, yielding a high-confidence regulatory network. For each gene *g*, we compute its total regulatory connectivity as:

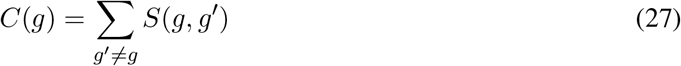

Genes with high total connectivity represent hub genes that broadly influence or are influenced by other CHD-associated genes.

##### Pathway Enrichment Analysis of the Response Vectors

To characterize the biological processes affected by each cluster of CHD genes, we perform pathway enrichment analysis on the genes most strongly affected by the knockouts. For each cluster *c*, we compute the mean response vector by averaging the response vectors of all assigned CHD genes:

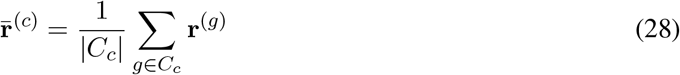

where *C*_*c*_ denotes the set of CHD genes in cluster *c*. We identify the top 200 most affected genes by selecting those with the lowest values in 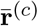, representing genes whose expression is most strongly suppressed by knockouts in this cluster. For each GOBP pathway from MSigDB, we test for enrichment in the top 200 genes using Fisher’s exact test (one-sided, greater alternative). Pathways with *p <* 0.05 are ranked by their *p*-values, and we report the top 20 for each cluster.

To investigate whether different CHD phenotypes are associated with disruption of distinct developmental programs, we partition CHD genes by their phenotype classification in the CHDgene database. For each phenotype, we compare pathway disruption patterns between phenotype-associated genes and non-associated genes. Specifically, for each gene, we identify the top 200 most affected genes from its knockout response vector and perform pathway enrichment analysis using Fisher’s exact test. For each GOBP pathway, we compute the fraction of genes whose knockouts significantly enrich that pathway (*p <* 0.05) separately for the phenotype-associated and non-associated groups. We calculate the enrichment ratio difference as the associated fraction minus the non-associated fraction. Pathways are ranked by this difference, where positive values indicate pathways preferentially disrupted by phenotype-associated genes. We report the top 20 pathways for each CHD phenotype.

##### Distinguishing ASD and VSD effects

To investigate whether Navigo captures chamber-specific gene expression programs, we analyze the impact of CHD gene knockouts on atrial versus ventricular marker genes. Using differential expression analysis between atrial and ventricular cardiomyocytes from the training data, we identify the top 50 atrial-enriched and top 50 ventricular-enriched marker genes. We partition CHD genes into four phenotype categories: ASD only, VSD only, both ASD and VSD, and other phenotypes. For each CHD gene knockout, we compute the mean absolute expression change for atrial and ventricular markers separately, then calculate the ventricular-to-atrial marker ratio. This ratio quantifies the relative impact on ventricular versus atrial gene expression programs. We compare these ratios across phenotype categories to determine whether ASD-associated genes preferentially affect atrial markers while VSD-associated genes preferentially affect ventricular markers.

### Evaluation of Genetic Perturbation Prediction

#### Experimental Setup

We evaluate Navigo’s ability to predict genetic knockout outcomes in a zero-shot manner using an independent large-scale scRNA-seq dataset profiling genetic perturbations during mouse embryogenesis [29]. This dataset was generated in a separate study from the training atlas, providing an independent test of model generalization. Mutants were generated through conventional gene-editing tools and breeding or tetraploid aggregation and collected at embryonic stage E13.5. Both wild-type and mutant embryos were profiled using sci-RNA-seq3. The dataset encompasses six genetic knockout perturbations: *Atp6v0a2, Gorab, Gli2, Ttc21b, Carm1*, and *Scn10a/11a* double knockout, with matched wild-type controls. To ensure compatibility with the training atlas, we retained only genes present in both datasets and applied consistent preprocessing. Cell type annotations were harmonized through manual mapping (details in Supplementary Table 1).

For each target gene *g* and cell type *c*, we compute the predicted gene expression change using the perturbation simulation approach described previously. We set the perturbation hyperparameter *m* = − 5. The predicted change for each gene *g*^′^ in cell type *c* is computed by averaging the difference between perturbed and wild-type trajectories across all cells within that type:

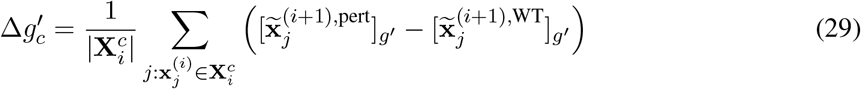

where 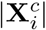 denotes the number of cells of type *c* at time point *t*_*i*_.

We train separate models for each major developmental trajectory, as different trajectories are governed by distinct gene regulatory networks with fundamentally different regulatory logic. This approach also mitigates the severe cell type imbalance issues across trajectories. For knockout predictions in a given cell type, we apply the vector field model corresponding to its major trajectory.

#### Metrics and Analysis

##### Evaluation of Prediction Accuracy

We rank genes by their predicted upregulation 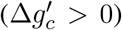 and downregulation 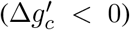. We compare the top 50 predictions in each direction against ground-truth DEGs identified from experimental knockout data using Wilcoxon rank-sum test with Benjamini-Hochberg correction (FDR *<* 0.05). Model performance is evaluated by the percentage of ground-truth DEGs recovered within the top 50 predictions.

##### Pathway Enrichment Analysis of the Predicted and Ground Truth Gene Expression Change

To evaluate whether predicted gene expression changes capture biologically meaningful perturbation responses, we performed pathway enrichment analysis comparing predicted and experimentally observed DEGs. For each knockout and cell type, we selected the top 100 upregulated genes from both predicted changes (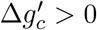, ranked by magnitude) and ground truth DEGs (FDR *<* 0.05). We tested enrichment against pathways relevant to the perturbed gene function from MSigDB v2025.1 (Mm) using the hyper-geometric test. We compared enrichment profiles between predicted and ground truth gene sets for each knockout and cell type.

##### Enrichment Analysis of the Downstream DEGs in Genetic Compensation

For each cell type, we simulated two perturbation scenarios: *Gli2* single knockout and *Gli1*/*Gli3* double knockout. To examine whether regulatory directionality among gene family members affects downstream gene sensitivity, we compared predicted expression changes between these scenarios. Genes were classified into two categories: (1) “Same Directions” genes, where predicted expression changes from both perturbations have the same sign (concordant responses); (2) “Different Directions” genes, where the perturbations induce opposite directional changes (discordant responses). For each category, we tested enrichment of experimentally validated *Gli2* knockout DEGs (FDR *<* 0.05) using the hypergeometric test. This analysis was performed independently for six cell types to assess lineage-specific patterns.

### Evaluation of Fate Reprogramming Prediction

#### Experimental Setup

We evaluate the ability of Navigo to identify TFs that can induce cell fate reprogramming from fibroblasts to various target cell types. Given fibroblasts as the source cell type *c*_*s*_ and a target cell type *c*_*t*_ (such as neurons, cardiomyocytes, or hepatocytes), we aim to predict which TF combinations can drive fibroblasts to adopt the transcriptomic state of *c*_*t*_. For each candidate TF set ℱ ⊆ {*f*_1_, *f*_2_, …, *f*_*n*_}, we compute the predicted gene expression change induced by TF overexpression using the perturbation simulation approach with hyperparameter *m* = 3:

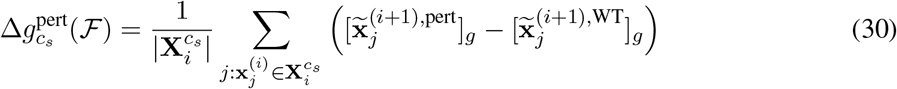

To evaluate whether TF set ℱ can drive fibroblasts toward target cell type *c*_*t*_, we compute the expected gene expression change as the difference between mean expression levels:

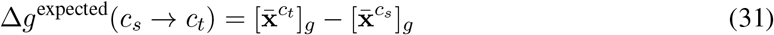

where 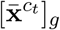 and 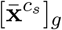 denote the mean expression of gene *g* in the target cell type and fibroblasts, respectively. Let *G*_top_(*c*_*s*_ → *c*_*t*_) denote the set of top DEGs between fibroblasts and the target cell type, and let *M* = |*G*_top_(*c*_*s*_ → *c*_*t*_)| . We then compute the directional alignment accuracy as the fraction of top DEGs whose predicted expression change matches the expected direction:

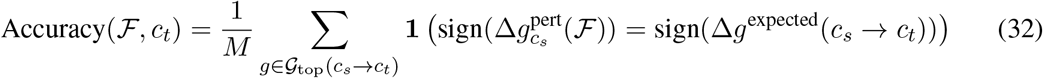

where **1**(·) is the indicator function. Higher alignment accuracy indicates stronger reprogramming potential. We rank candidate TF sets by their alignment accuracy to identify the most promising TF combinations for each target cell type.

We validated predictions using two independent sources of experimental evidence. The first comprised curated data from published literature [97, 98, 130, 131, 132, 133] documenting established reprogramming factors across multiple target cell types (Supplementary Figure 19a). The second was derived from a comprehensive experimental screen [31] that systematically evaluated hundreds of pairwise TF combinations from the bHLH and POU families for their ability to reprogram mouse fibroblasts into functional neurons (Supplementary Figure 19b). This screen identified 76 TF pairs that successfully induced neuronal differentiation, with quantitative characterization of reprogramming efficiency through morphological, electrophysiological, and transcriptomic analyses. This validation approach enabled both broad evaluation across diverse lineages and fine-grained assessment of combinatorial TF interactions.

We train the developmental vector field model using data from fibroblasts and the target cell type of interest. This focused training strategy enables the model to learn the specific regulatory dynamics governing transitions between these two cell states.

#### Metrics and Analysis

##### AUROC and AUPRC Metrics for TF ranking across lineages

For each target cell type *c*_*t*_, we defined the positive set *P* (*c*_*t*_) as all validated TF combinations for and the negative set 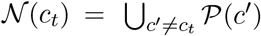 as all TF combinations validated for other cell types. For each candidate TF set ℱ ∈ *P* (*c*_*t*_) ∪ *N* (*c*_*t*_), we computed Accuracy(ℱ, *c*_*t*_). We then ranked all candidate sets by their accuracy scores and evaluated prediction performance using AUROC and AUPRC, treating sets in *P*(*c*_*t*_) as positive labels and sets in *N*(*c*_*t*_) as negative labels. To ensure robustness, we trained three model checkpoints and generated five bootstrap samples (80% cell subsampling) per target cell type, reporting mean metrics across evaluations.

##### Metrics for Combinatorial Ranking across two TF families

For bHLH-POU combinatorial screening, we leveraged experimental data profiling neuronal reprogramming efficiency for TF pairs (*f*_*i*_, *f*_*j*_) where *f*_*i*_ ∈ ℱ_bHLH_ and *f*_*j*_ ∈ ℱ_POU_. Let *E*(*f*_*i*_, *f*_*j*_) denote experimental efficiency and *S*(*f*_*i*_, *f*_*j*_) = Accuracy({*f*_*i*_, *f*_*j*_}) denote predicted accuracy. We classified pairs as positive labels if *E*(*f*_*i*_, *f*_*j*_) > 0 (successful reprogramming) and as negative labels if *E*(*f*_*i*_, *f*_*j*_) = 0 (failed reprogramming), then evaluated AUROC using *S* as the ranking score. To reduce noise, we applied two filters: (1) programming effect threshold PE_thr, setting *E*(*f*_*i*_, *f*_*j*_) → 0 if *E*(*f*_*i*_, *f*_*j*_) < PE_thr; (2) expression threshold Expr_thr, excluding TFs *f* with mean expression < Expr_thr. We systematically evaluated PE_thr ∈ {0, 0.2, 0.4, 0.6, 0.8, 1.0} and Expr_thr ∈ {0, 0.005, 0.01, 0.02} . For anchor-based screening, we fixed one TF *f* ^∗^ ∈ ℱ_family1_ and ranked all partners *f* ∈ ℱ_family2_ by *S*(*f* ^∗^, *f*), computing AUROC for each anchor. We examined the correlation between TF mean expression and anchor-based AUROC to assess whether prediction accuracy depends on expression levels.

## Data availability

All the datasets we use are listed in Methods and are publicly available. The mouse embryogenesis scRNA-seq datasets are available at https://omg.gs.washington.edu/jax/public/download.html (GSE186069 and GSE228590). The mouse mutant datasets are available at https://atlas.gs.washington.edu/mmca_v2/ (GSE199308).

## Code availability

Code for the package and reproducing the results is in the GitHub repository: https://github.com/aristoteleo/Navigo-release. The package introduction and tutorial notebooks are also available at https://navigo.readthedocs.io/en/latest/.

## Acknowledgements

This study was supported by multiple funding sources, including the Chinese University of Hong Kong (CUHK; award numbers 4937025, 4937026, 5501517, 5501329 and SHIAE BME-p1-24 to Y.L.), the IdeaBooster Fund (IDBF23ENG05 and IDBF24ENG06 to Y.L.), a grant from the Research Grants Council of the Hong Kong Special Administrative Region (HKSAR), China (project numbers CUHK 24204023 and 14208525 to Y.L.), and grants from the Innovation and Technology Commission of the HKSAR, China (project numbers GHP/065/21SZ, ITS/247/23FP and PRP/033/24FX to Y.L.). This research was also partially supported by the Research Matching Grant Scheme at CUHK (award numbers 8601603 and 8601663 to Y.L.) from the Research Grants Council, HKSAR, China. X. Q. acknowledged funding support from the Laude Moonshot Seed Grant, Pantas And Ting Sutardja Foundation, the Wu Tsai Neurosciences Institute Big Ideas in Neuroscience Program.

## Author contributions

Y.L. and X.Q. conceived and supervised the research. Y.F. implemented the models. Y.F., Y.L. and X.Q. designed the downstream experiments. Y.F. and X.L. ran the experiments, interpreted and visualized the results. Y.F., X.L. and Y.W. wrote the manuscript. All authors read and approved the final manuscript.

