## Supplementary Materials for "Generative Modeling of Mouse Embryogenesis for Fate and Disease Prediction"

[1]Department of Computer Science and Engineering, CUHK, Hong Kong SAR, China
[2]Department of Genetics, Stanford University School of Medicine, Stanford, CA, USA
[3]Basic Sciences and Engineering Initiative, Betty Irene Moore Children's Heart Center, Lucile Packard Children's Hospital, Stanford, CA, USA
[4]Department of Computer Science, Stanford University, Stanford, CA, USA
[5]Stanford Cardiovascular Institute, Stanford University, Stanford, CA, USA
[6]The CUHK Shenzhen Research Institute, Hi-Tech Park, Nanshan, Shenzhen, China
[7]Institute for Medical Engineering and Science, Massachusetts Institute of Technology, Cambridge, MA, USA
[8]Wyss Institute for Biologically Inspired Engineering, Harvard University, Boston, MA, USA
[9]Broad Institute of MIT and Harvard, Cambridge, MA, USA
[10]Current Address: Canchen Technology, Shenzhen, China
[*]These authors contributed equally to this work

#### List of Tables

1 Cell type mapping between the KO dataset and the training dataset. For one-to-many mappings, the corresponding cell types in the training dataset are separated by commas. . . . . . . . . . . 25

22 (**Related to Figure 5**) Relationship between expression level and prediction accuracy for POU TFs. 23

### A Supplementary Figures

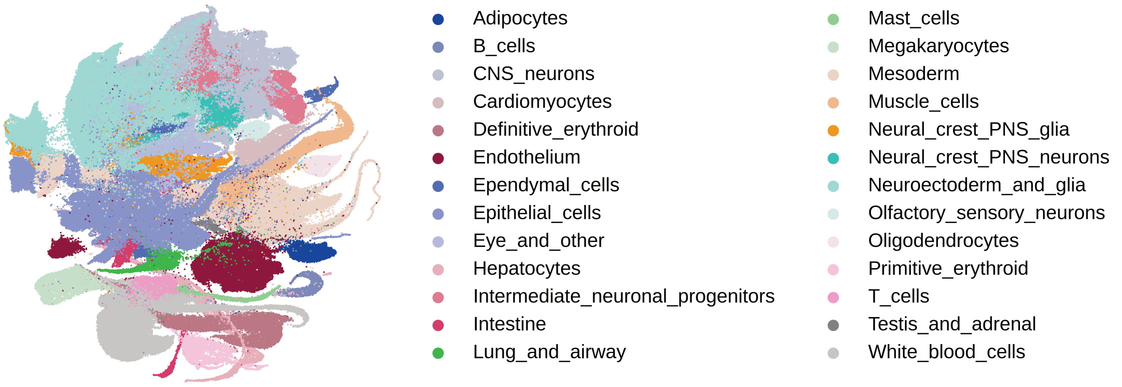

Supplementary Figure 1: (**Related to Figure 1**) UMAP visualization of the dataset colored by major developmental trajectories

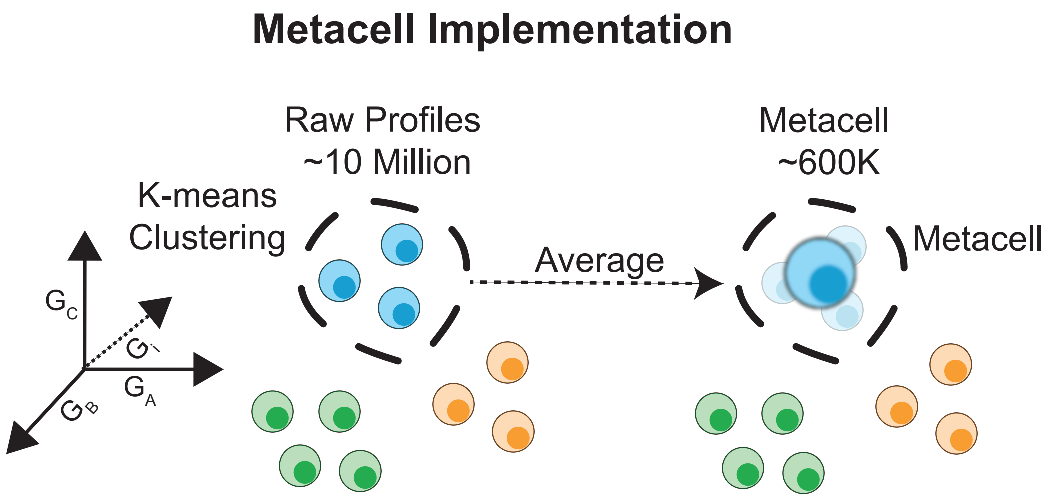

Supplementary Figure 2: (**Related to Figure 1**) Illustration of the implementation of Metacell.

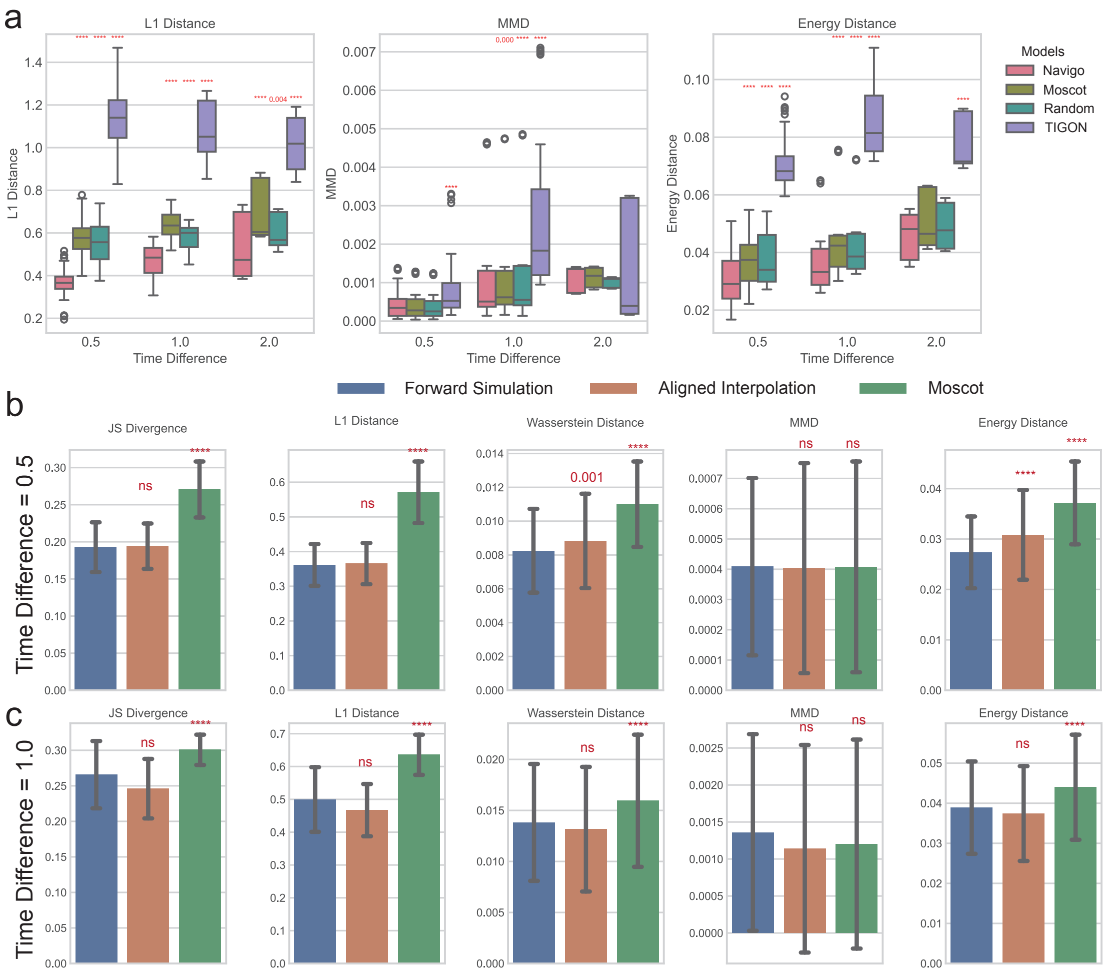

Supplementary Figure 3: (**Related to Figure 2**) Comparison of interpolation performance between Navigo and baseline models. **a**. L1 distance, MMD and Energy Distance of Navigo and baseline models, evaluated on three settings with different time intervals. The two-tailed paired Wilcoxon test results are annotated with red text. $n$=95,40,15 for interval=0.5,1,2. **b, c**. Performance comparison of two interpolation modes, forward simulation and aligned interpolation, and Moscot in two settings with different time intervals. Two-sided Wilcoxon tests are performed to test if forward simulation performs significantly better than others. $p$-value is annotated as red text (****: $p$-val<0.0001, ns: $p$-val>0.05). For panel b, $n$=95, for panel c, $n$=40.

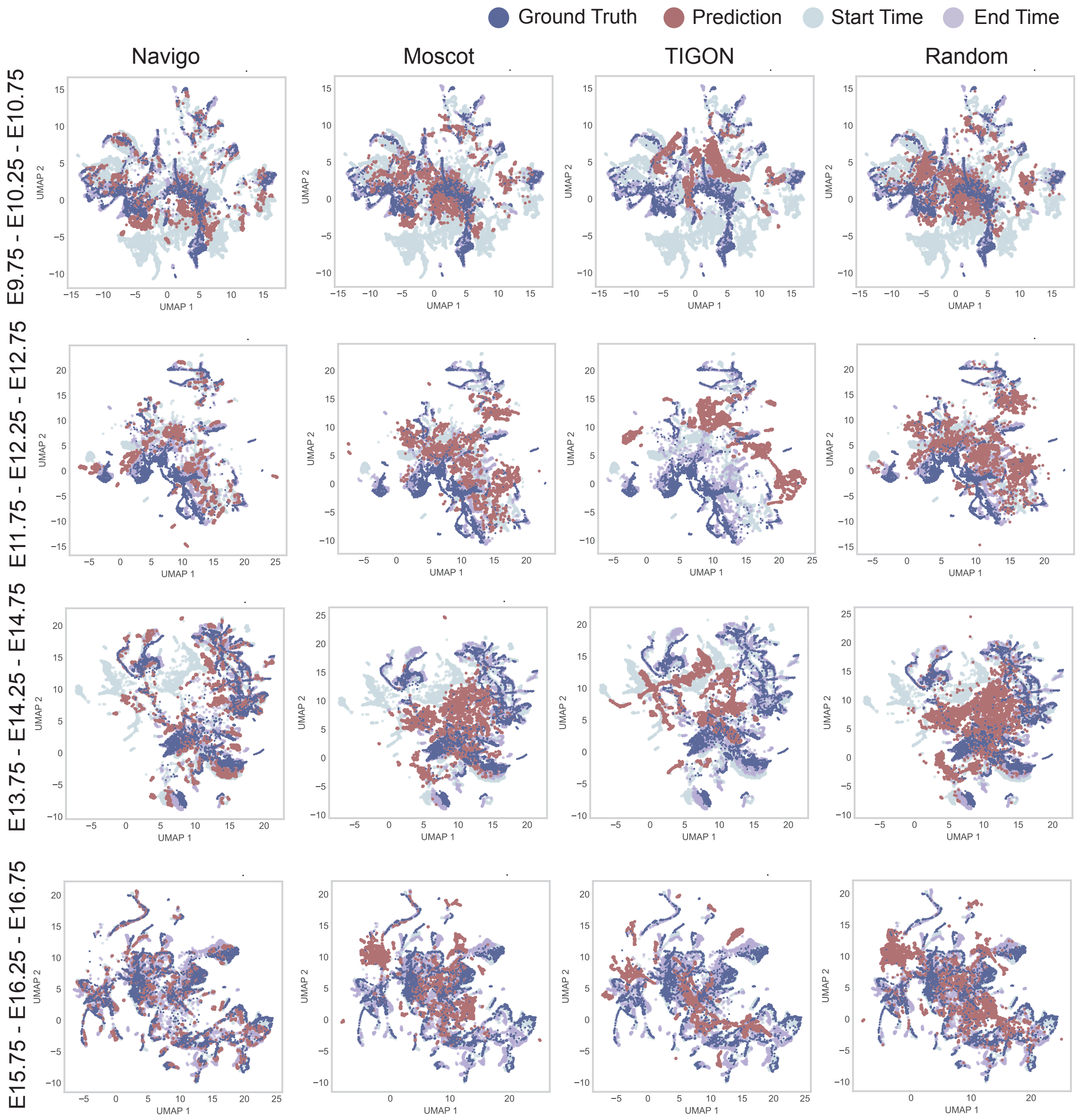

Supplementary Figure 4: (**Related to Figure 2**) UMAP visualization of interpolation results. The predicted and ground truth gene expression profiles are shown in red and blue points, respectively. Four cases with different interpolation time points are visualized.

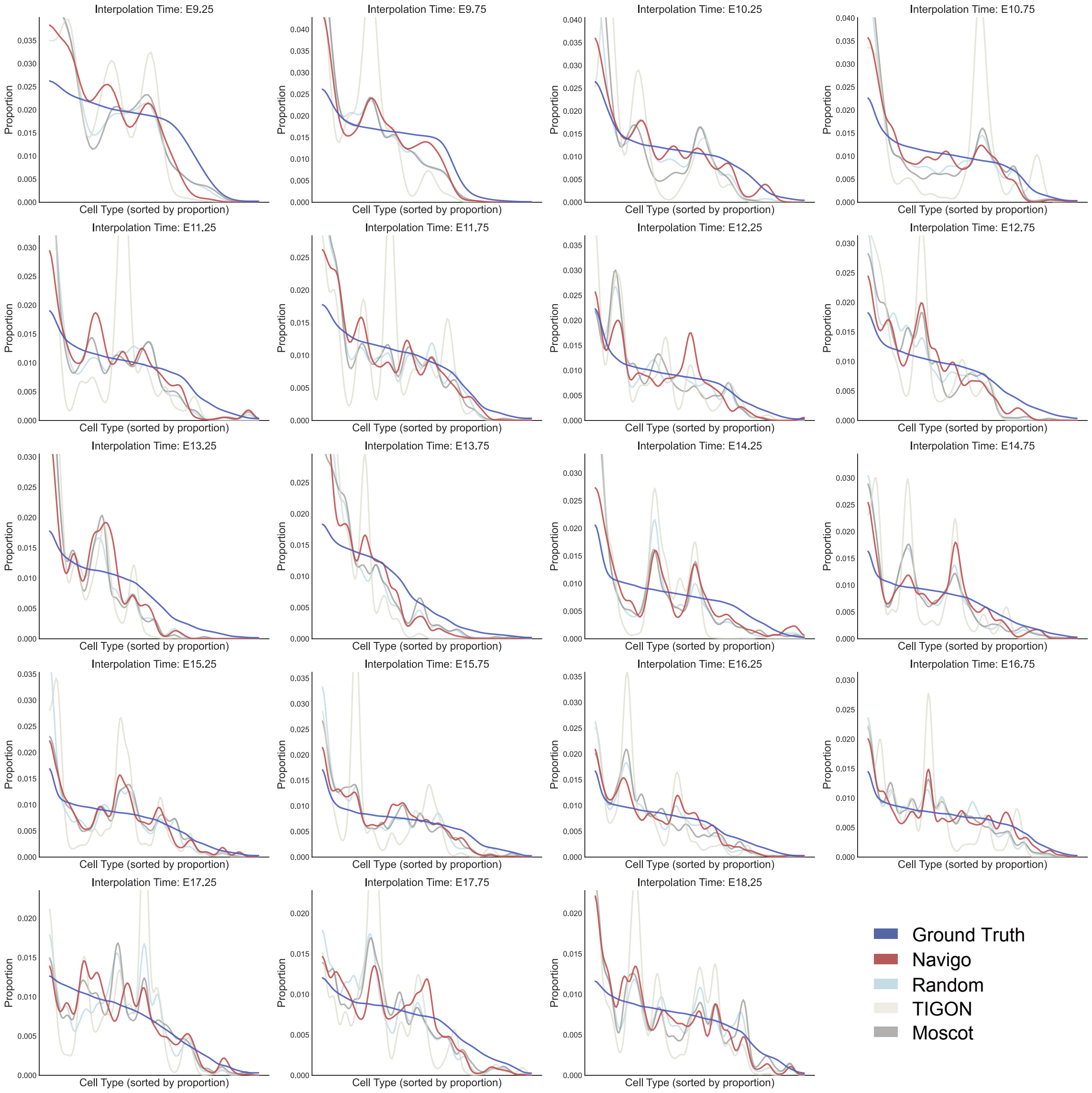

Supplementary Figure 5: (**Related to Figure 2**) Predicted cell type proportion distribution curves of different interpolation time points. The time interval is 0.5. Ground truth distribution curves are shown as blue lines.

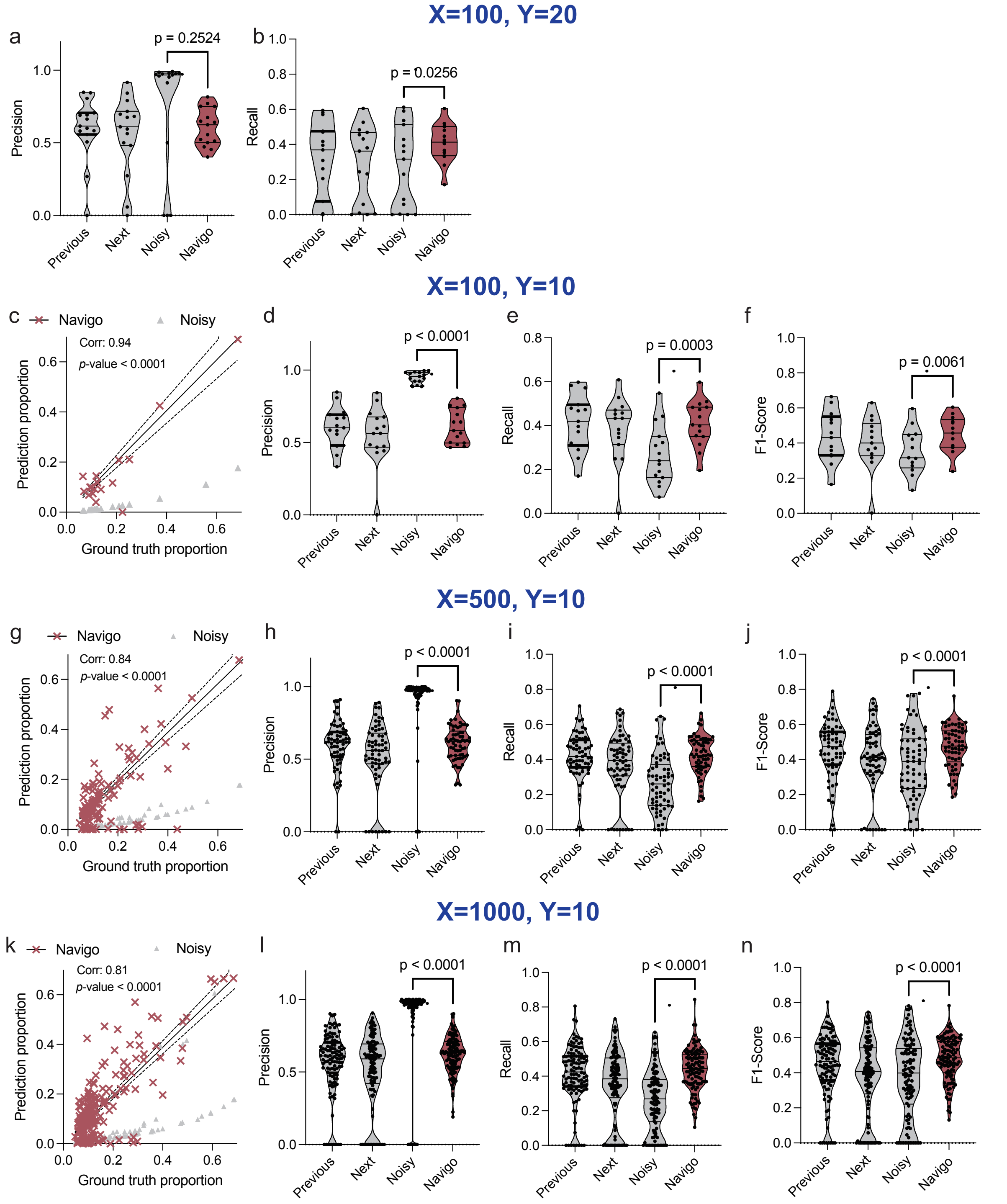

Supplementary Figure 6: (**Related to Figure 2**) Denoising performance on simulated data under different hyperparameter settings. For $X = 100$ and $Y = 20$, **a** and **b** show marker gene prediction precision and recall, respectively. For $X = 100$ and $Y = 10$, **c–f** show the cell type proportion prediction scatter plot, marker gene prediction precision, recall, and F1-score, respectively ($n$=15). For $X = 500$ and $Y = 10$, **g–j** show the same metrics in the same order ($n$=65). For $X = 1000$ and $Y = 10$, **k–n** show the same metrics in the same order ($n$=110).

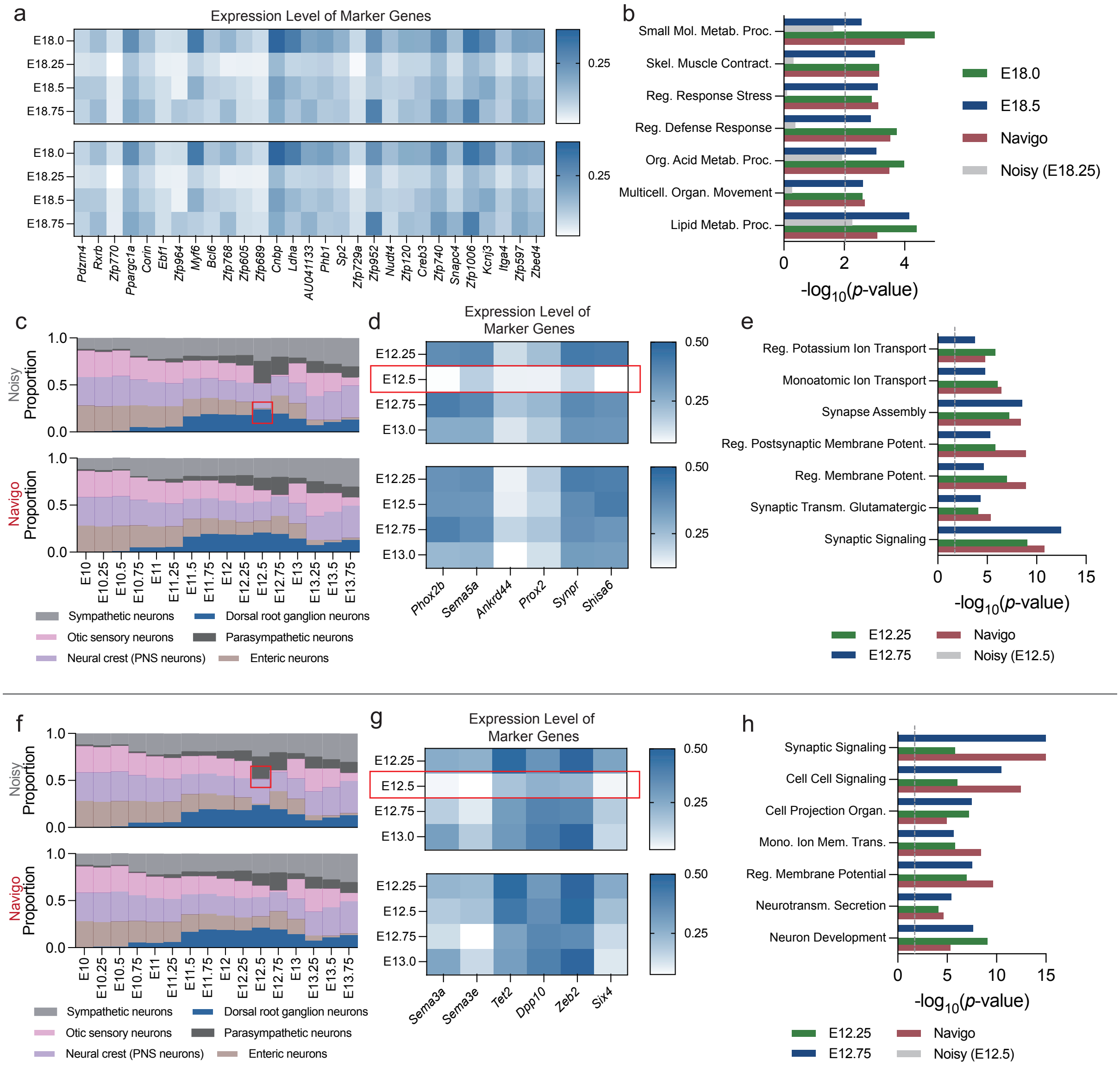

Supplementary Figure 7: (**Related to Figure 2**) Denoising performance on real data with additional case studies. **a**. Extension of Figure 2j in the main text. **b.** Enriched biological pathways in original, predicted, and neighboring myofibroblast populations at E18.25. **c**. Comparison of original and Navigo-predicted cell type proportions along the neural crest PNS neurons developmental trajectory across time points. Enteric neuron proportions at E12.5 exhibit apparent noise in the original data. **d**. Marker gene expression dynamics along the Enteric neuron developmental trajectory in original versus predicted cell populations. **e**. Enriched biological pathways in original, predicted, and neighboring Enteric neuron populations at E12.5. **f**. Comparison of original and Navigo-predicted cell type proportions along the neural crest PNS neurons developmental trajectory across time points. Otic sensory neuron proportions at E12.5 exhibit apparent noise in the original data. **g**. Marker gene expression dynamics along the otic sensory neuron developmental trajectory in original versus predicted cell populations. **h**. Enriched biological pathways in original, predicted, and neighboring otic sensory neuron populations at E12.5.

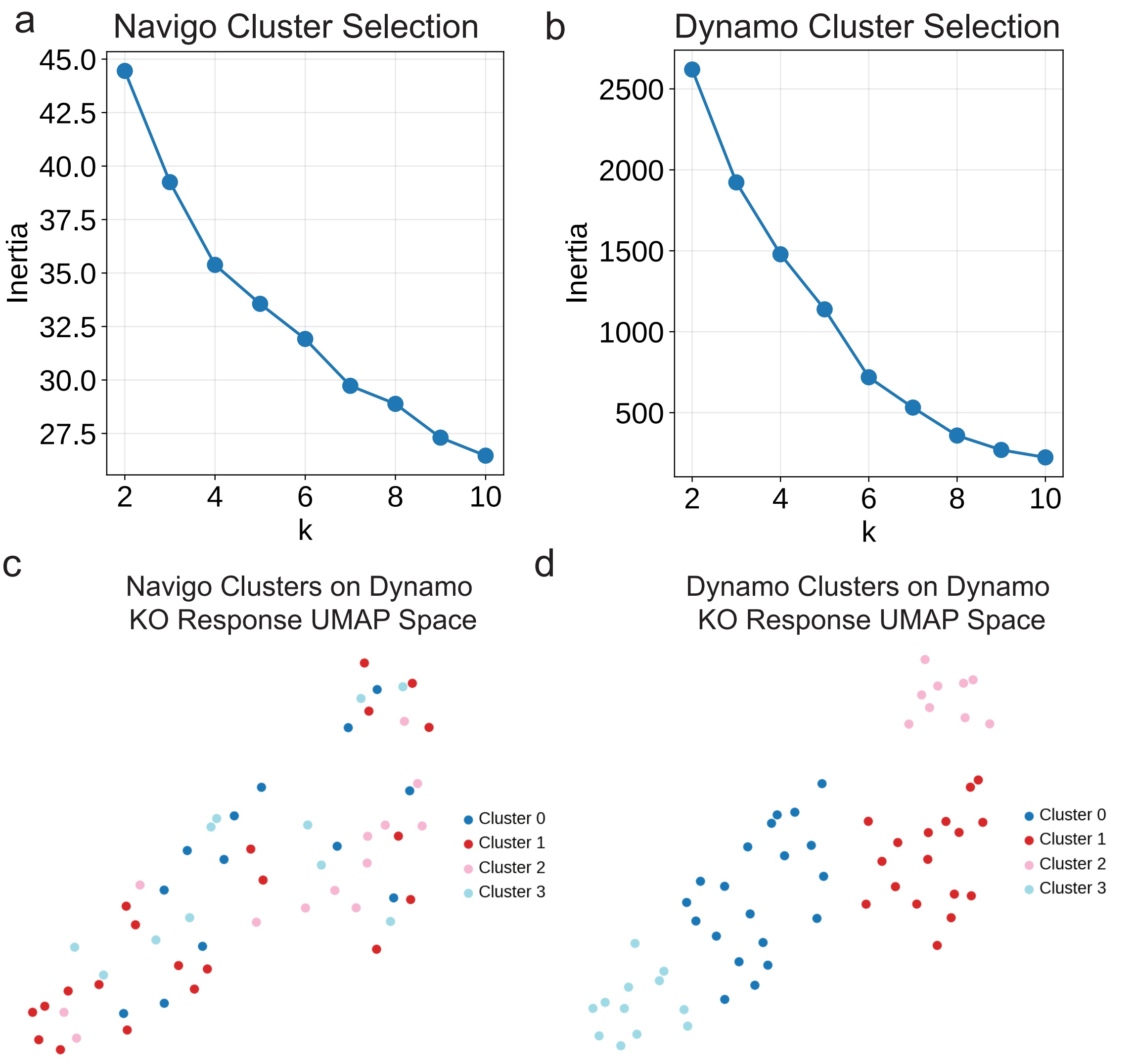

Supplementary Figure 8: (**Related to Figure 3**) Hyperparameter selection and performance comparison for clustering CHD gene responses. **a**. Inertia plot for clustering of Navigo gene knockout response vectors. **b**. Inertia plot for clustering of Dynamo gene knockout response vectors. **c**. UMAP visualization of the Dynamo gene knockout response vectors colored by cluster labels obtained from the Navigo vectors. **d**. UMAP visualization of the Dynamo gene knockout response vectors colored by cluster labels obtained from the Dynamo vectors.

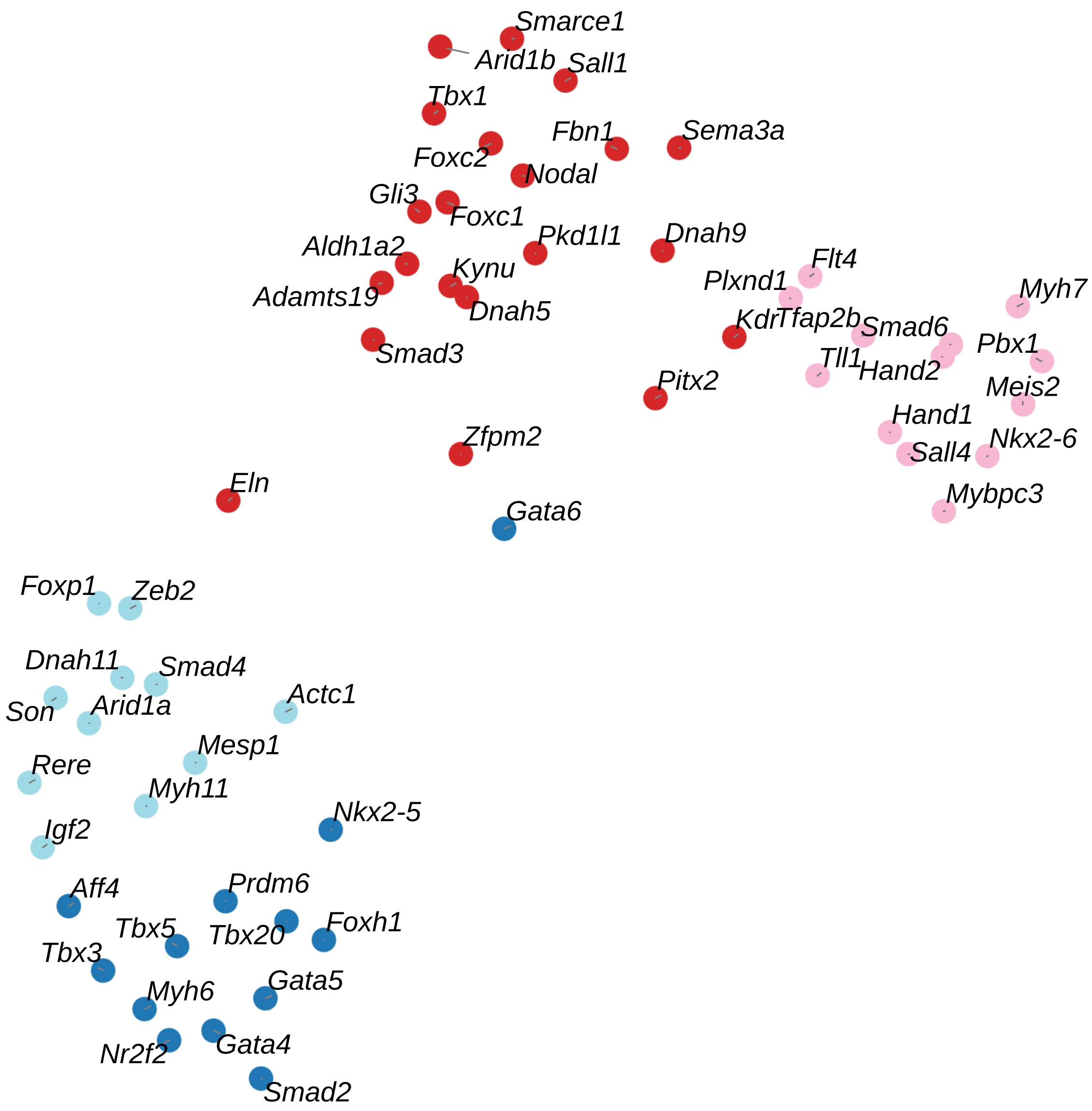

Supplementary Figure 9: (**Related to Figure 3**) UMAP visualization of gene knockout response vectors reveals functional clustering. Each point represents the transcriptomic response to knocking out a specific CHD-associated gene, annotated with the gene name. Clusters correspond to distinct developmental processes.

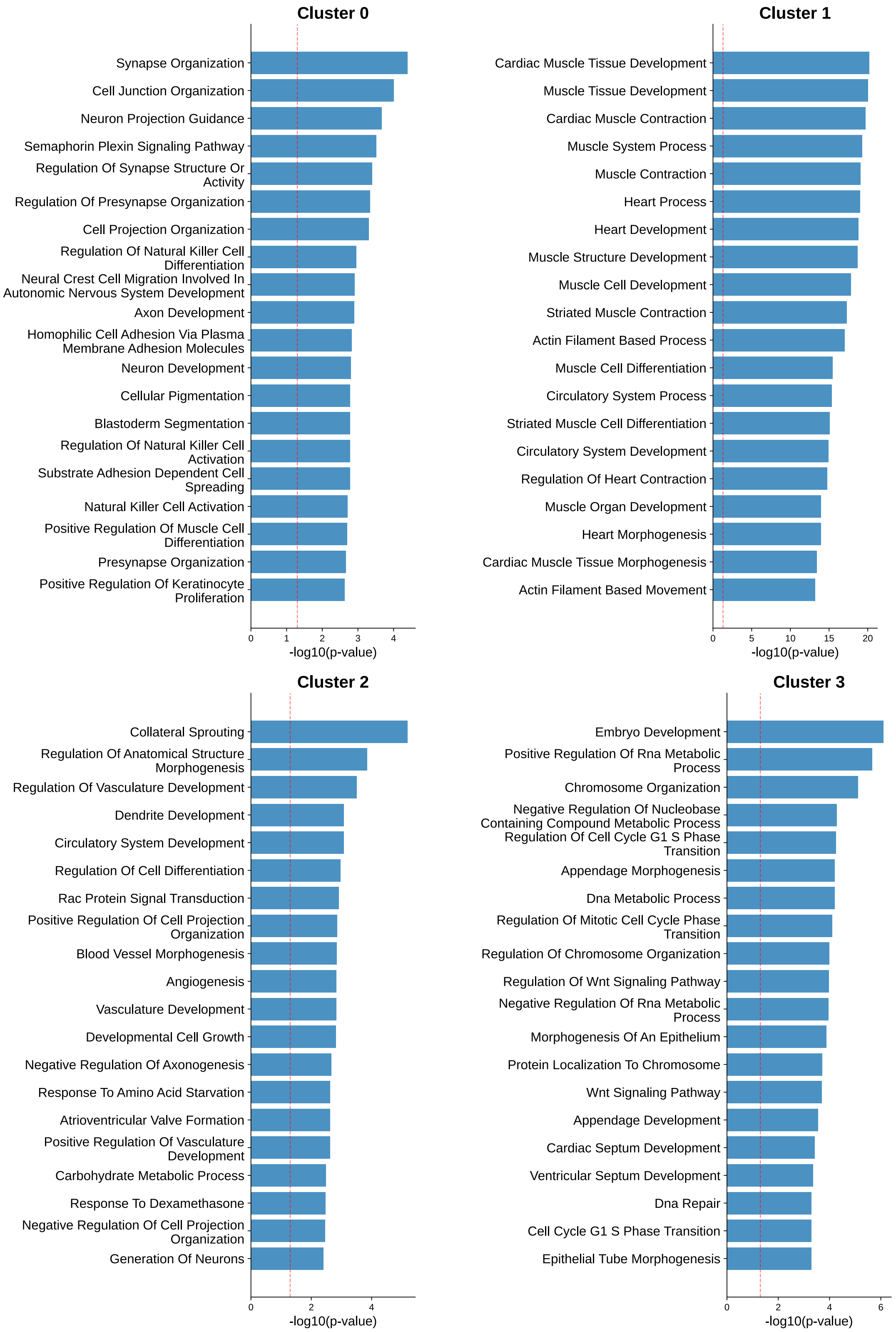

Supplementary Figure 10: (**Related to Figure 3**) GOBP pathway enrichment analysis for each cluster in the Navigo CHD gene clustering results. Hypergeometric tests are conducted on the top affected genes in the average response vector for each cluster.

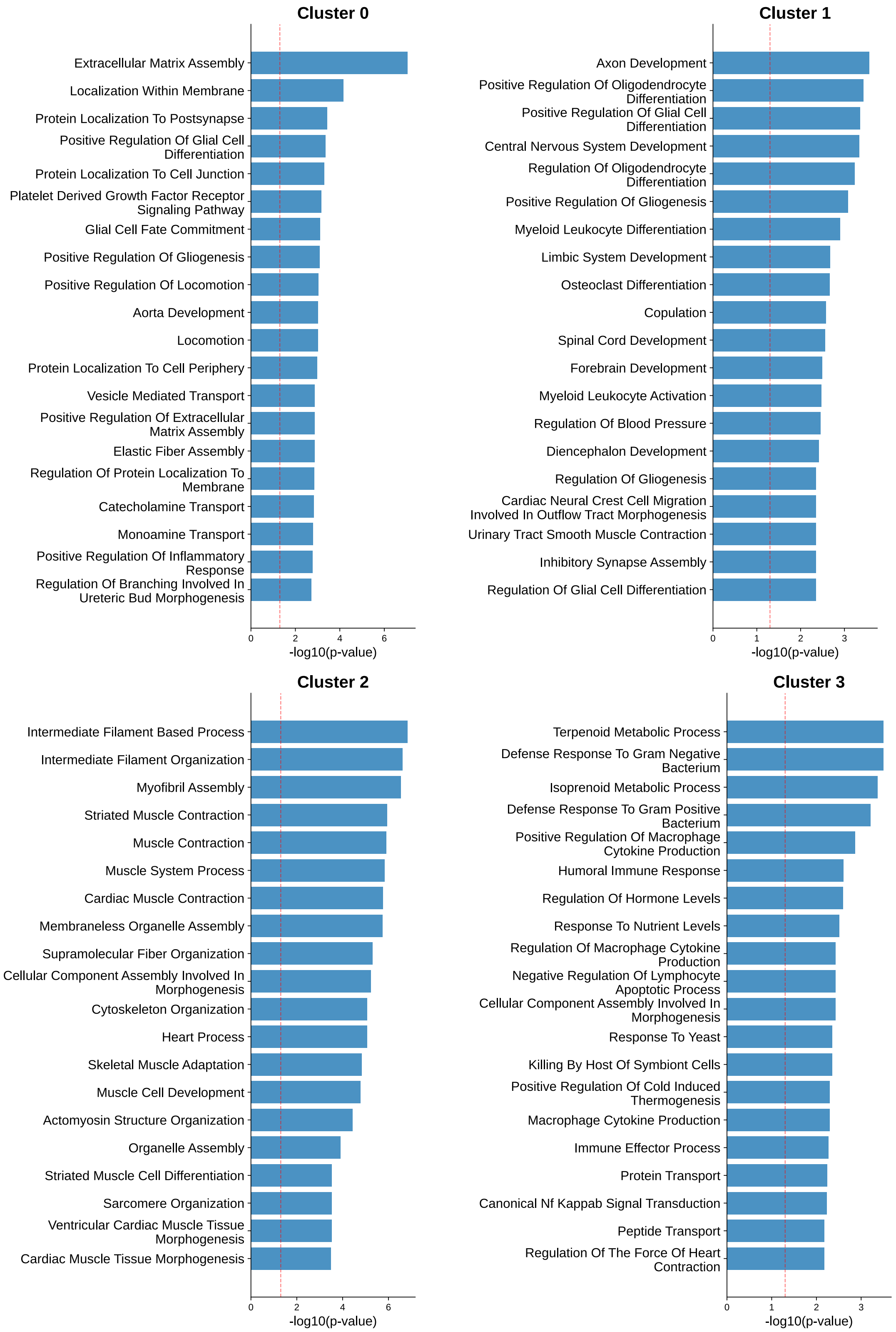

Supplementary Figure 11: (**Related to Figure 3**) GOBP pathway enrichment analysis for each cluster in the Dynamo CHD gene clustering results. Hypergeometric tests are conducted on the top affected genes in the average response vector for each cluster.

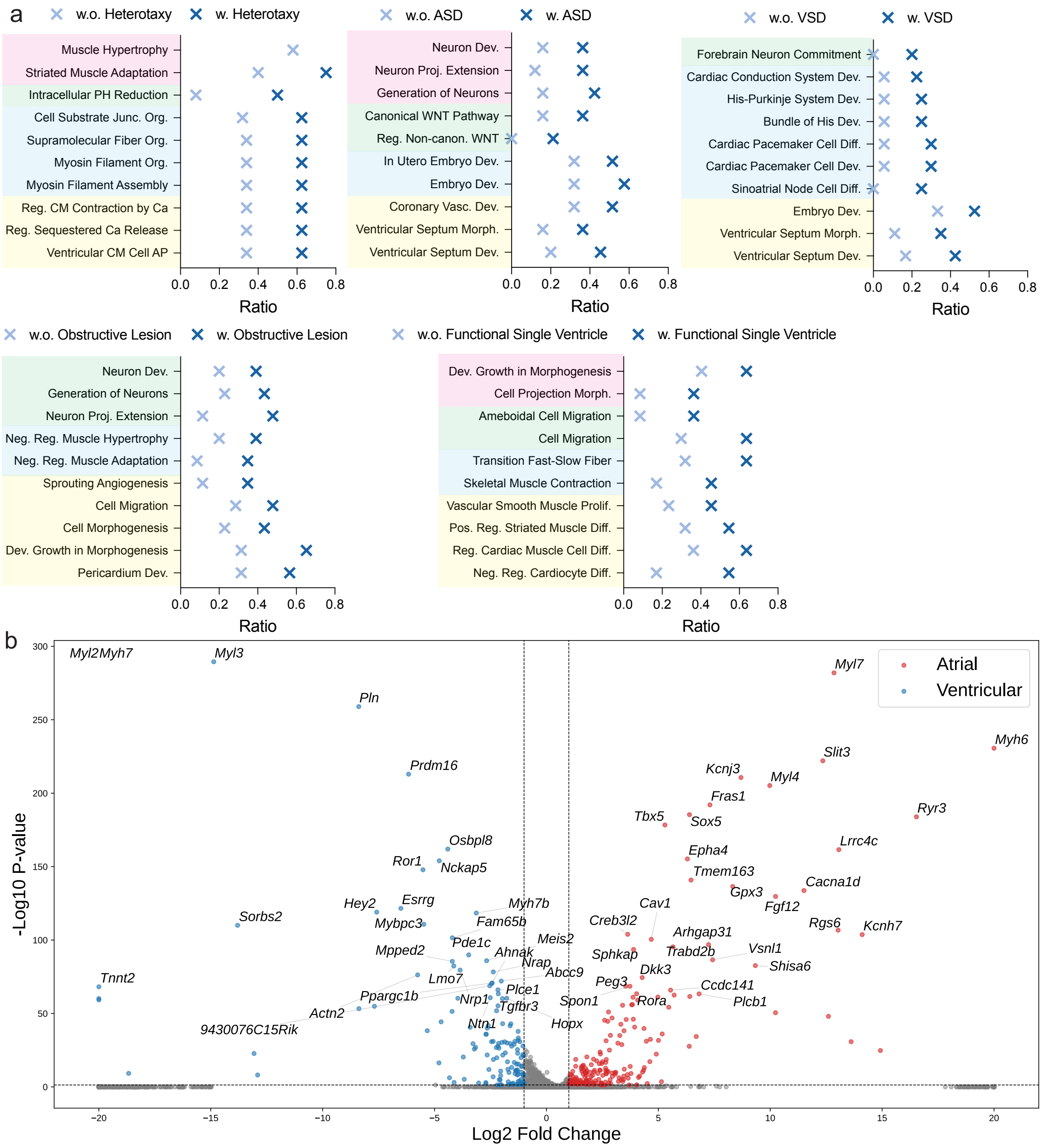

Supplementary Figure 12: (**Related to Figure 3**) Disease pathway enrichment patterns for all CHD subtypes and DEGs between atrial and ventricular cardiomyocytes. **a**. The top affected pathways in each CHD subtype. The pathways are manually clustered based on their functions. We used hypergeometric tests without correction. **b**. The volcano plot for the differentially expressed genes (DEGs) between atrial and ventricular cardiomyocytes. We used Wilcoxon rank-sum tests implemented with scanpy for calculation.

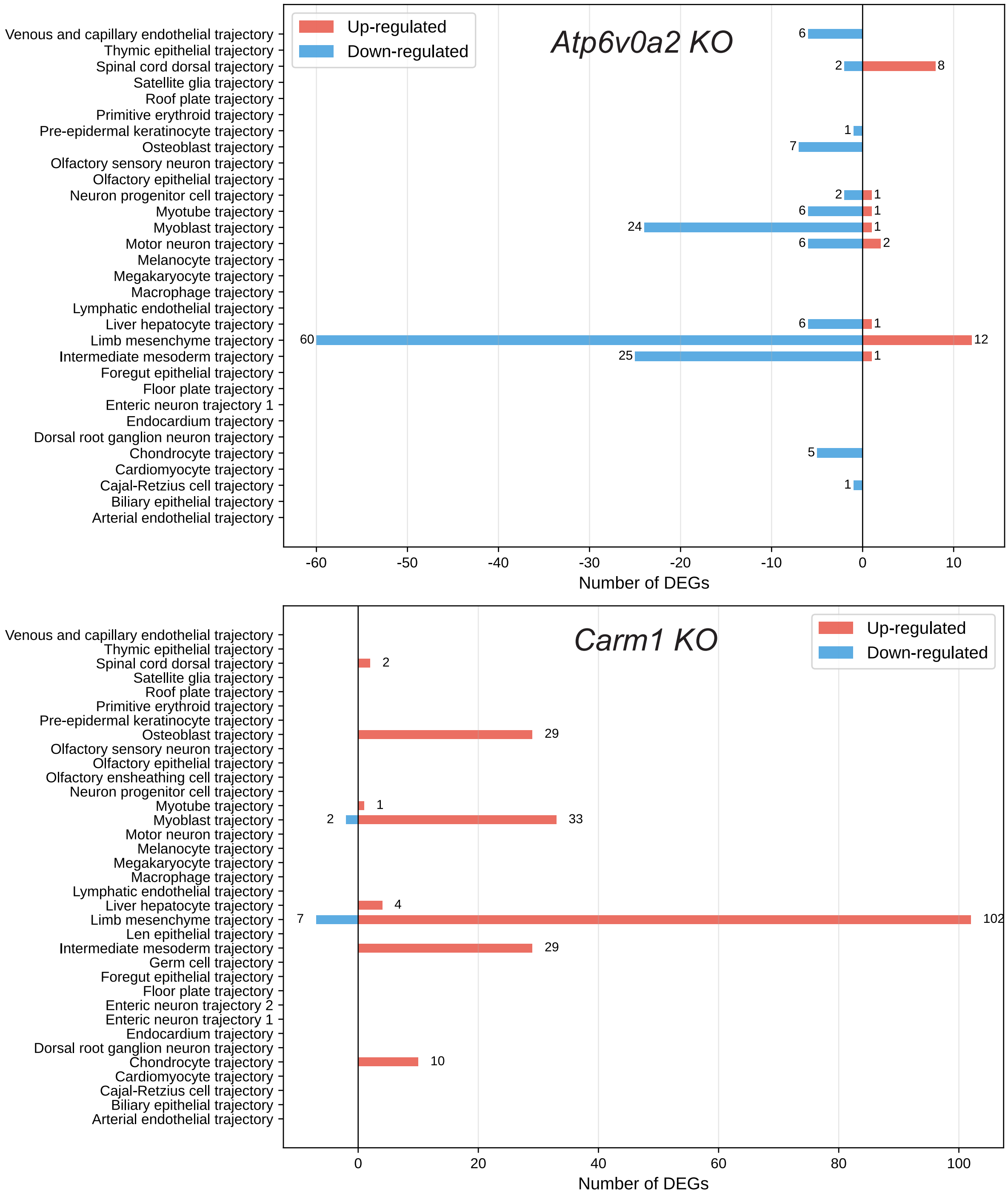

Supplementary Figure 13: (**Related to Figure 4**) Statistics of the number of up and down DEGs across different cell types with respect to *Atp6v0a2* KO and *Carm1* KO

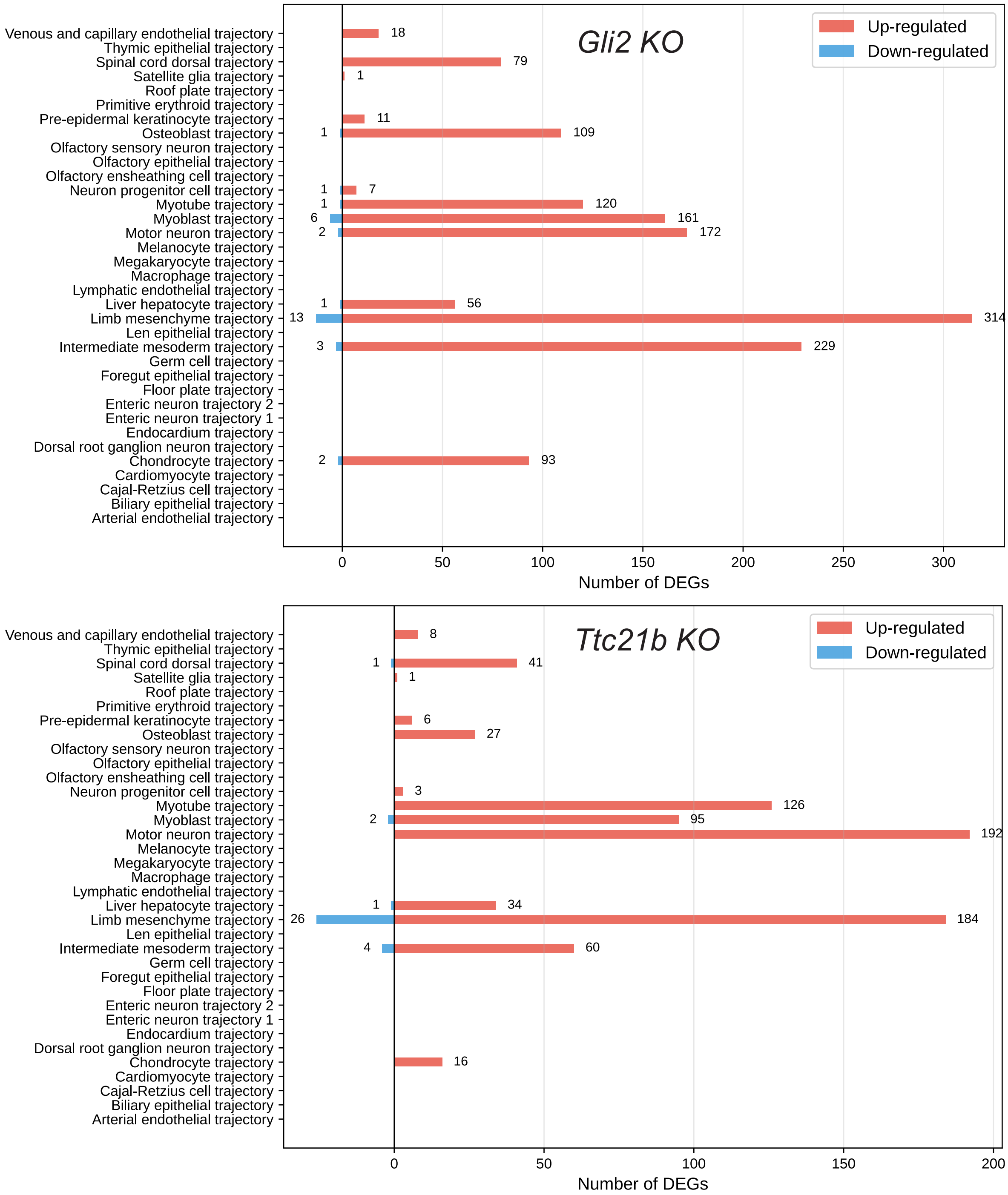

Supplementary Figure 14: (**Related to Figure 4**) Statistics of the number of up and down DEGs across different cell types with respect to *Gli2* KO and *Ttc21b* KO

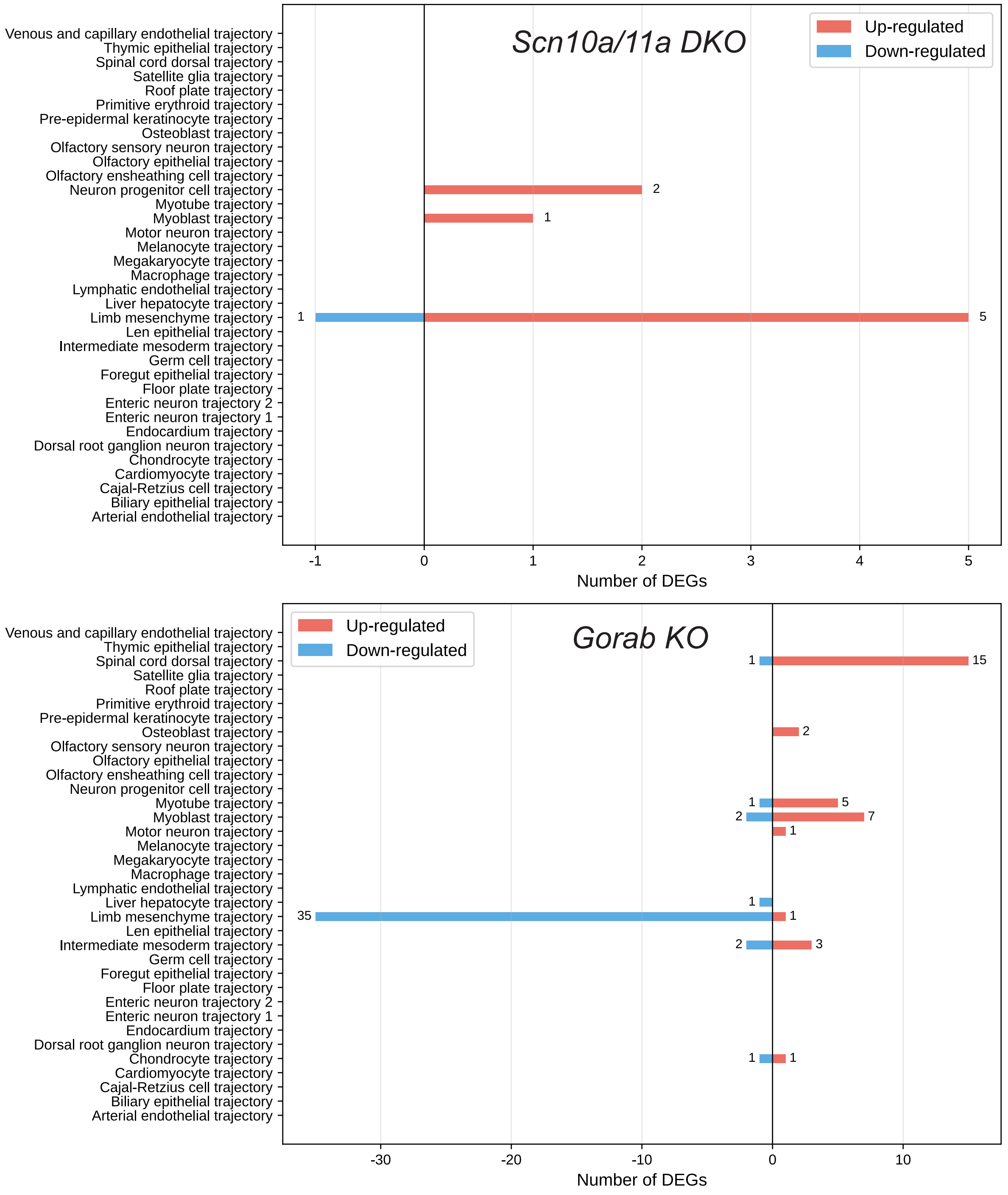

Supplementary Figure 15: (**Related to Figure 4**) Statistics of the number of up and down DEGs across different cell types with respect to *Scn10a/11a* DKO and *Gorab* KO

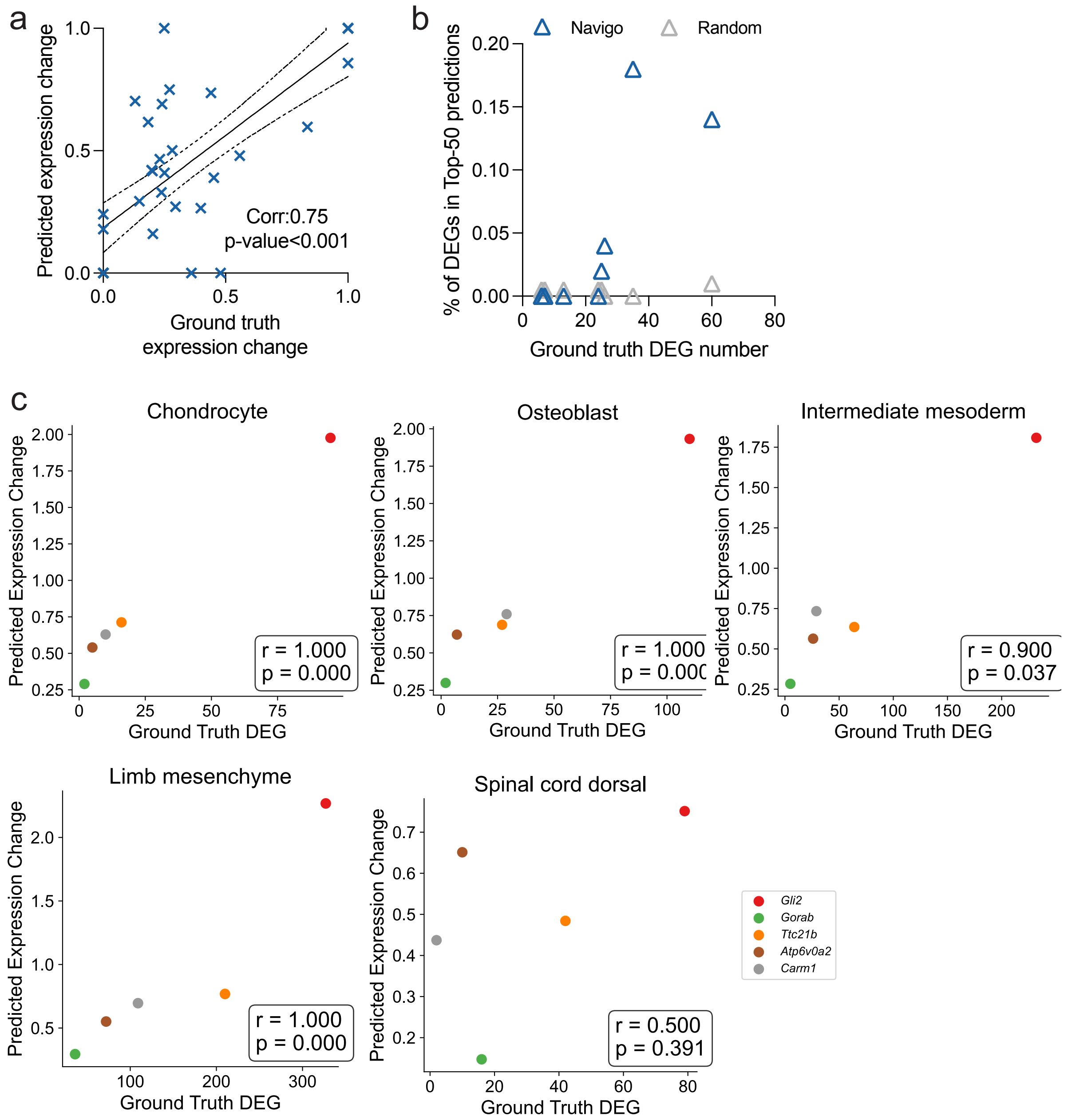

Supplementary Figure 16: (**Related to Figure 4**) Supplementary benchmarks for Figure 4(c-e) in the manuscript. **a**. Correlation between the mean absolute ground truth expression change and predicted expression changes across different gene KO and cell type pairs. $n$=35. **b**. Relationship between knockout effect size, defined by the number of ground truth down-regulated DEGs, and prediction accuracy. **c**. The relationship between the predicted expression change after gene KO and the number of ground truth DEGs across different lineages. Each dot indicates one gene KO. The Spearman rank-order correlation coefficient is reported, with the $p$-value shown. $n$=5.

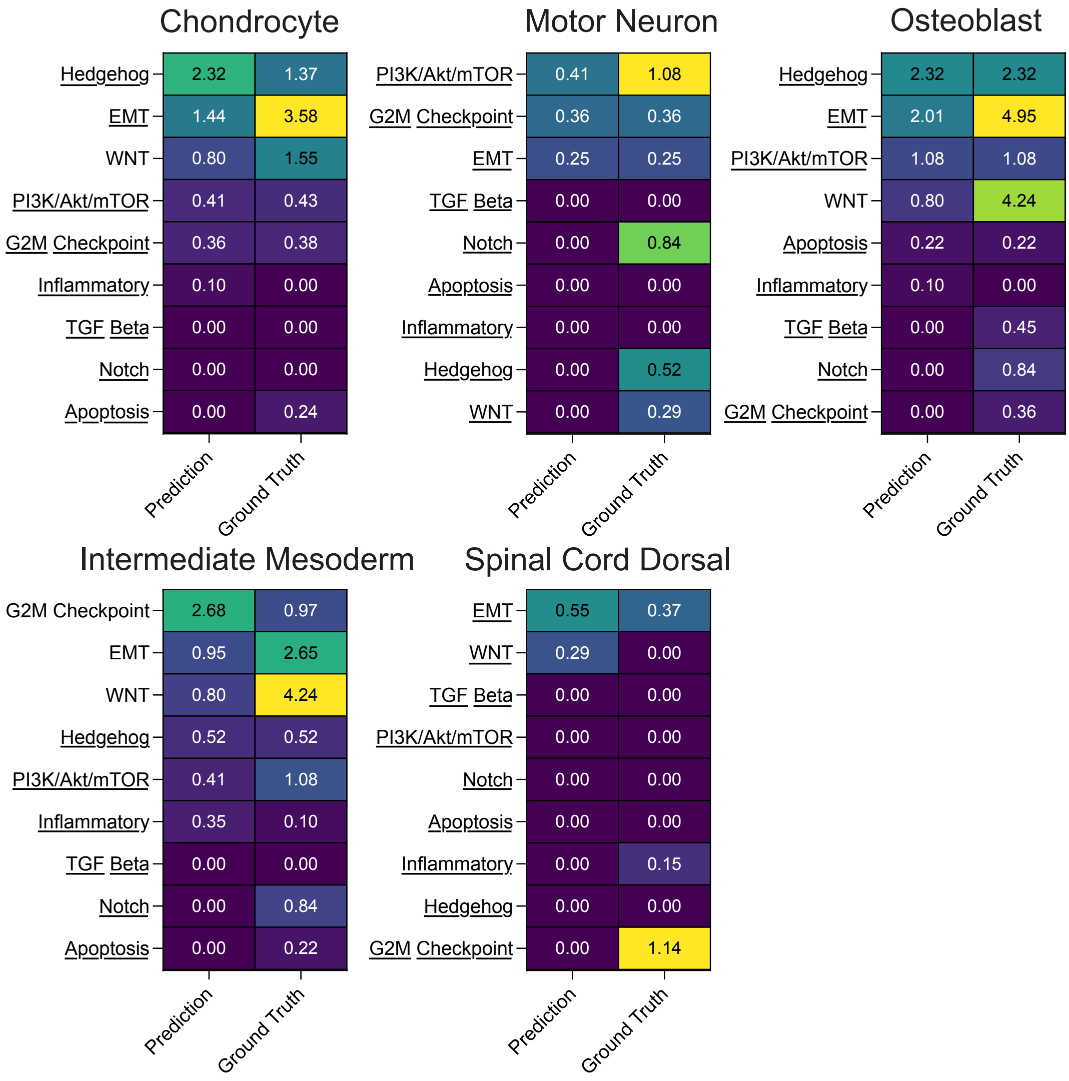

Supplementary Figure 17: (**Related to Figure 4**) Comparison of enriched pathways between predicted top up-regulated genes and ground-truth DEGs across lineages. The heatmap shows $-\log_{10}(p\text{-value})$ from hypergeometric tests for pathway enrichment in predicted top up-regulated genes (left columns) and ground-truth DEGs (right columns). The significance threshold is $-\log_{10}(0.05) \approx 1.3$. Underlined pathways indicate alignment between predicted and ground-truth enrichment (both significant or both insignificant).

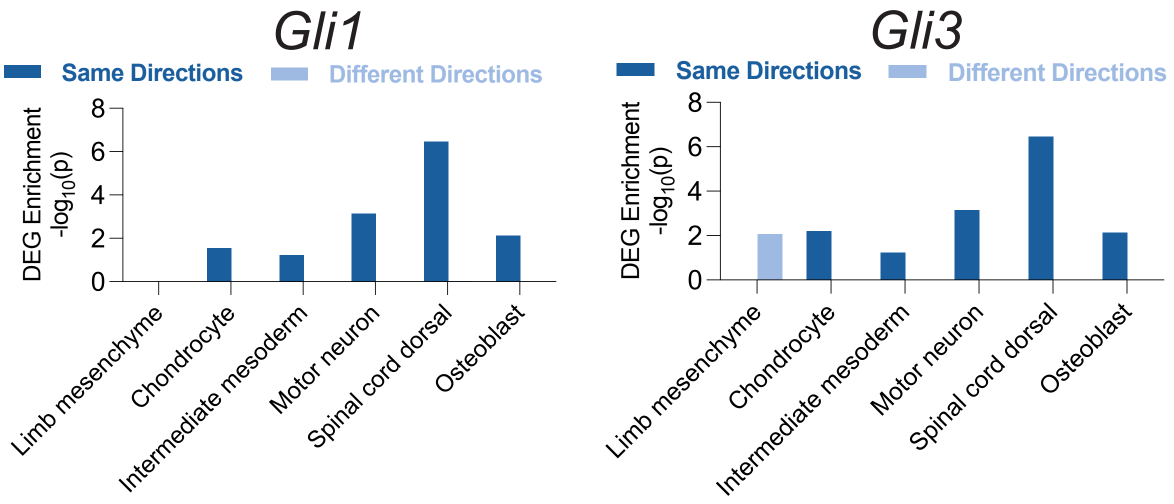

Supplementary Figure 18: (**Related to Figure 4**) Regulatory directionality predicts gene sensitivity to *Gli1* and *Gli3* knockout. Enrichment of experimental *Gli1* KO and *Gli3* KO DEGs in "Same Directions" versus "Different Directions" categories across lineages. The directional categories are defined by comparing simulated *Gli1* KO with *Gli2* KO, and simulated *Gli3* KO with *Gli2* KO, respectively. We used hypergeometric tests without correction.

a

| Source Cells | Target Cells | Key TFs |
|---|---|---|
| Fibroblast | Cardiomyocyte | *GATA4, MEF2C, TBX5* |
| Fibroblast | Skeletal muscle cell | *MyoD1* |
| Fibroblast | Neurons | *ASCL1, POU3F2, MYT1L* |
| Fibroblast | Hepatocytes | *HNF4A, FOXA1, FOXA2, FOXA3* |
| Fibroblast | Pancreatic Beta Cells | *PDX1, NEUROG3, MAFA* |
| Fibroblast | Endothelial Cells | *ETV2, ERG, FLI1, FOXC2* |

b

| | *Ascl1* | *Ascl2* | *Ascl3* | *Ascl5* | *Neurog1* | *Neurog2* | *Neurog3* | *Neurod1* | *Neurod2* | *Neurod4* | *Neurod6* | *Atoh1* | *Atoh7* | *Atoh8* | *Myf5* | *Ptf1a* |
|---|---|---|---|---|---|---|---|---|---|---|---|---|---|---|---|---|
| *Pou4f3* | 5.70 | 3.00 | 0.02 | 0.95 | 0.42 | 0.29 | 2.00 | 0.16 | 0.31 | 0.00 | 0.02 | 0.30 | 0.04 | 0.00 | 0.05 | 0.00 |
| *Pou4f1* | 3.20 | 1.80 | 0.02 | 0.29 | 0.33 | 0.32 | 0.82 | 0.03 | 0.63 | 0.06 | 0.02 | 0.26 | 0.11 | 0.01 | 0.00 | 0.00 |
| *Pou4f2* | 2.80 | 2.70 | 0.00 | 0.00 | 0.21 | 0.28 | 0.97 | 0.20 | 0.33 | 0.00 | 0.00 | 0.05 | 0.00 | 0.00 | 0.04 | 0.01 |
| *Pou3f2* | 1.60 | 1.40 | 0.00 | 0.00 | 0.76 | 0.56 | 1.80 | 0.07 | 0.02 | 0.00 | 0.00 | 0.01 | 0.00 | 0.00 | 0.00 | 0.00 |
| *Pou3f4* | 0.20 | 0.15 | 0.00 | 0.00 | 0.36 | 0.13 | 3.00 | 0.04 | 0.00 | 0.00 | 0.00 | 0.00 | 0.00 | 0.00 | 0.00 | 0.00 |
| *Pou3f3* | 0.00 | 0.00 | 0.00 | 0.00 | 0.66 | 0.09 | 0.15 | 0.02 | 0.00 | 0.00 | 0.00 | 0.00 | 0.00 | 0.00 | 0.00 | 0.00 |
| *Pou3f1* | 0.03 | 0.00 | 0.00 | 0.00 | 0.13 | 0.02 | 0.11 | 0.01 | 0.00 | 0.00 | 0.00 | 0.00 | 0.00 | 0.00 | 0.00 | 0.00 |
| *Pou1f1* | 0.18 | 0.25 | 0.00 | 0.00 | 0.02 | 0.03 | 0.34 | 0.06 | 0.00 | 0.00 | 0.00 | 0.00 | 0.00 | 0.00 | 0.00 | 0.00 |
| *Pou5f1* | 1.10 | 0.55 | 0.00 | 0.00 | 0.06 | 0.05 | 0.84 | 0.07 | 0.01 | 0.00 | 0.00 | 0.02 | 0.00 | 0.00 | 0.00 | 0.00 |
| *Nr4a2* | 0.05 | 0.03 | 0.00 | 0.00 | 0.00 | 0.00 | 0.00 | 0.00 | 0.00 | 0.00 | 0.00 | 0.00 | 0.00 | 0.00 | 0.00 | 0.00 |

Supplementary Figure 19: (**Related to Figure 5**) Statistics for the datasets used in fibroblast reprogramming prediction. **a**. Key TFs for reprogramming fibroblasts into various cell types, compiled from established literature. **b**. Ground truth efficiency heatmap showing reprogramming outcomes for combinations of bHLH and POU family transcription factors in fibroblast-to-neuron reprogramming.

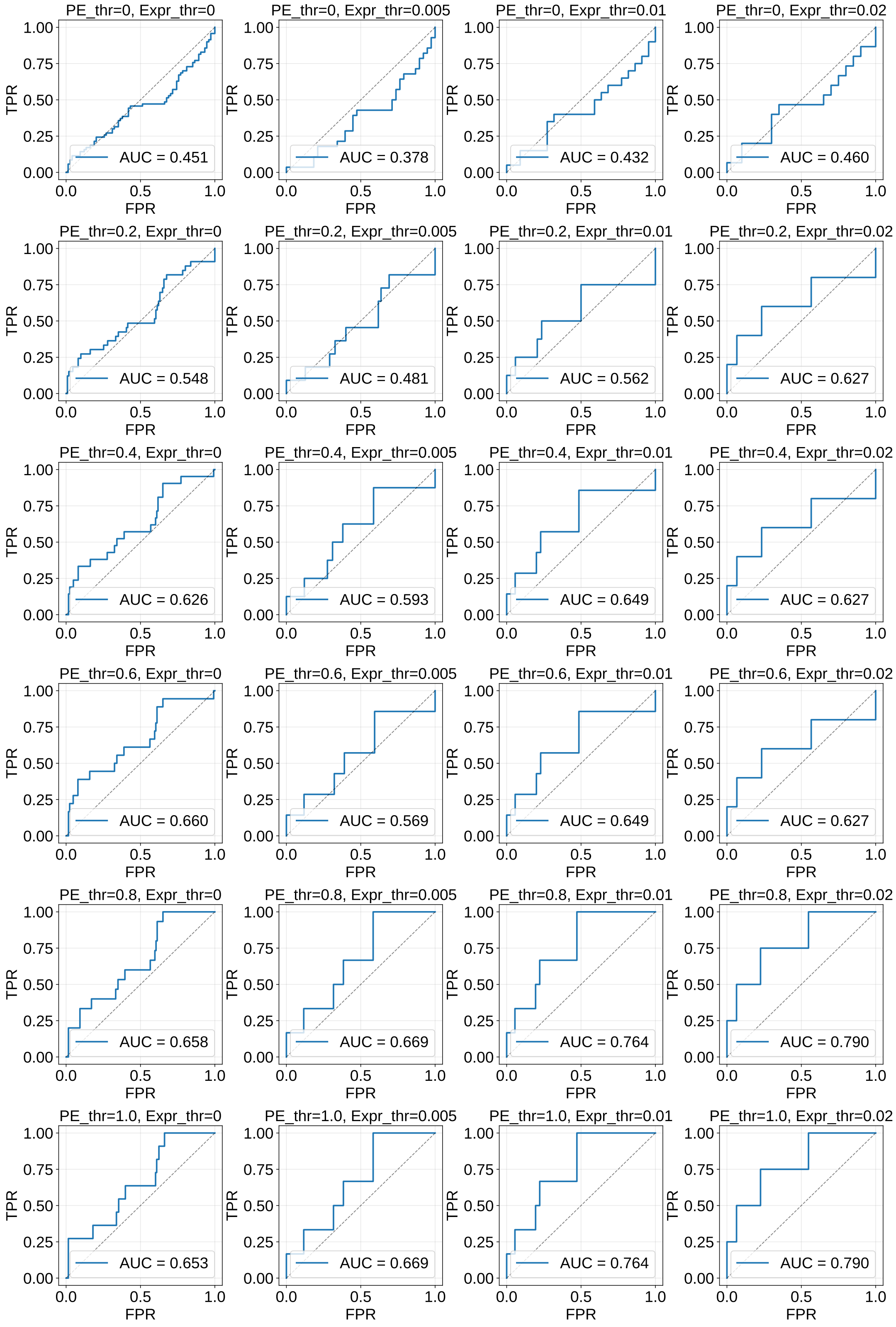

Supplementary Figure 20: (**Related to Figure 5**) The screening performance measured by AUROC for bHLH and POU transcription factors. We systematically varied two hyperparameters: the programming effect threshold, which treats experimentally measured programming effects below a specified value as zero, and the expression threshold, which restricts modeling to transcription factors with average expression levels above a defined cutoff.

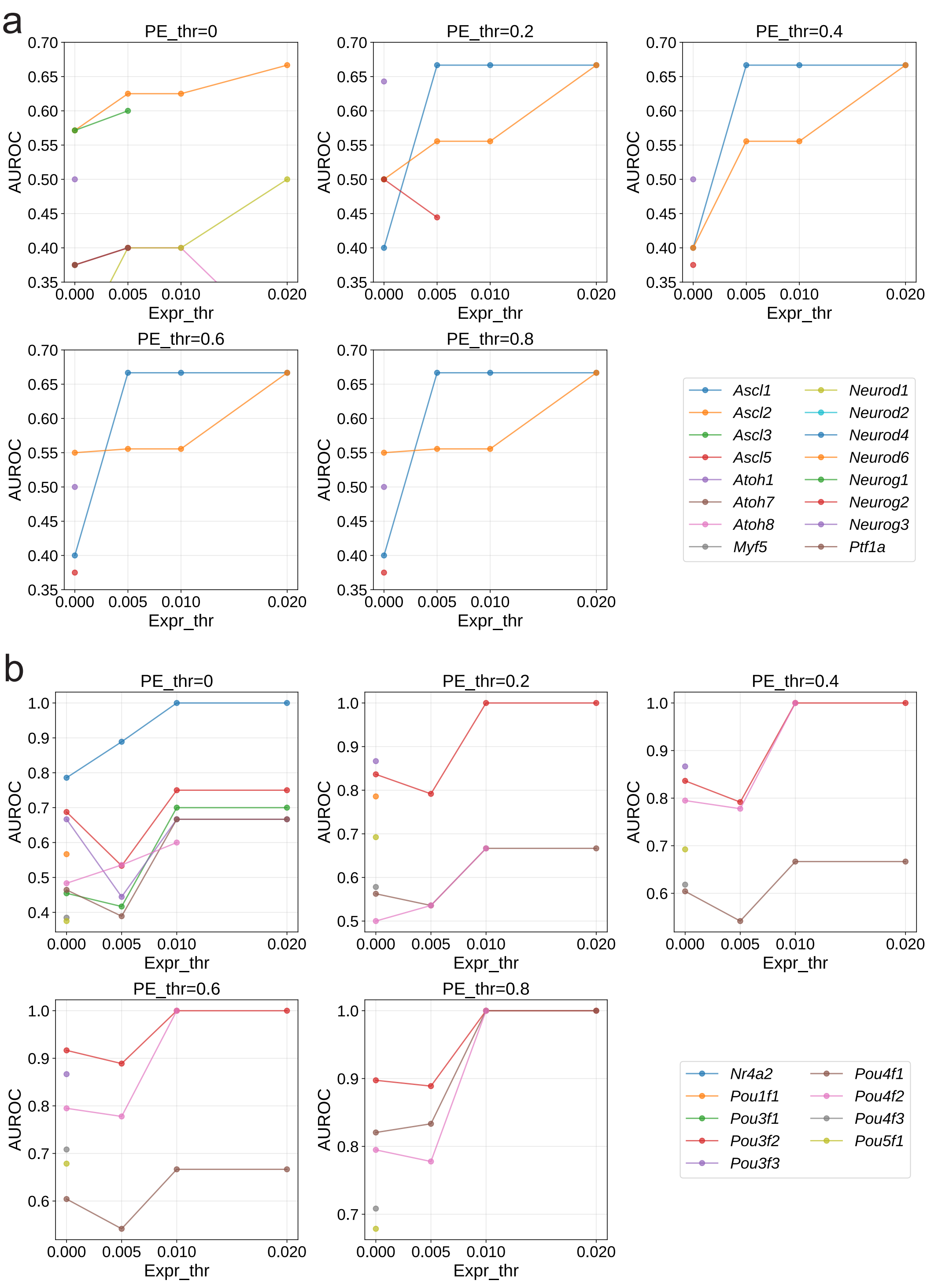

Supplementary Figure 21: (**Related to Figure 5**) Impact of expression threshold and programming effect threshold on anchor-based TF screening performance. **a**. Each POU family member was fixed as an anchor while screening bHLH TFs as partners. AUROC measures prediction accuracy for each POU anchor. **b**. Each bHLH family member was fixed as an anchor while screening POU TFs as partners.

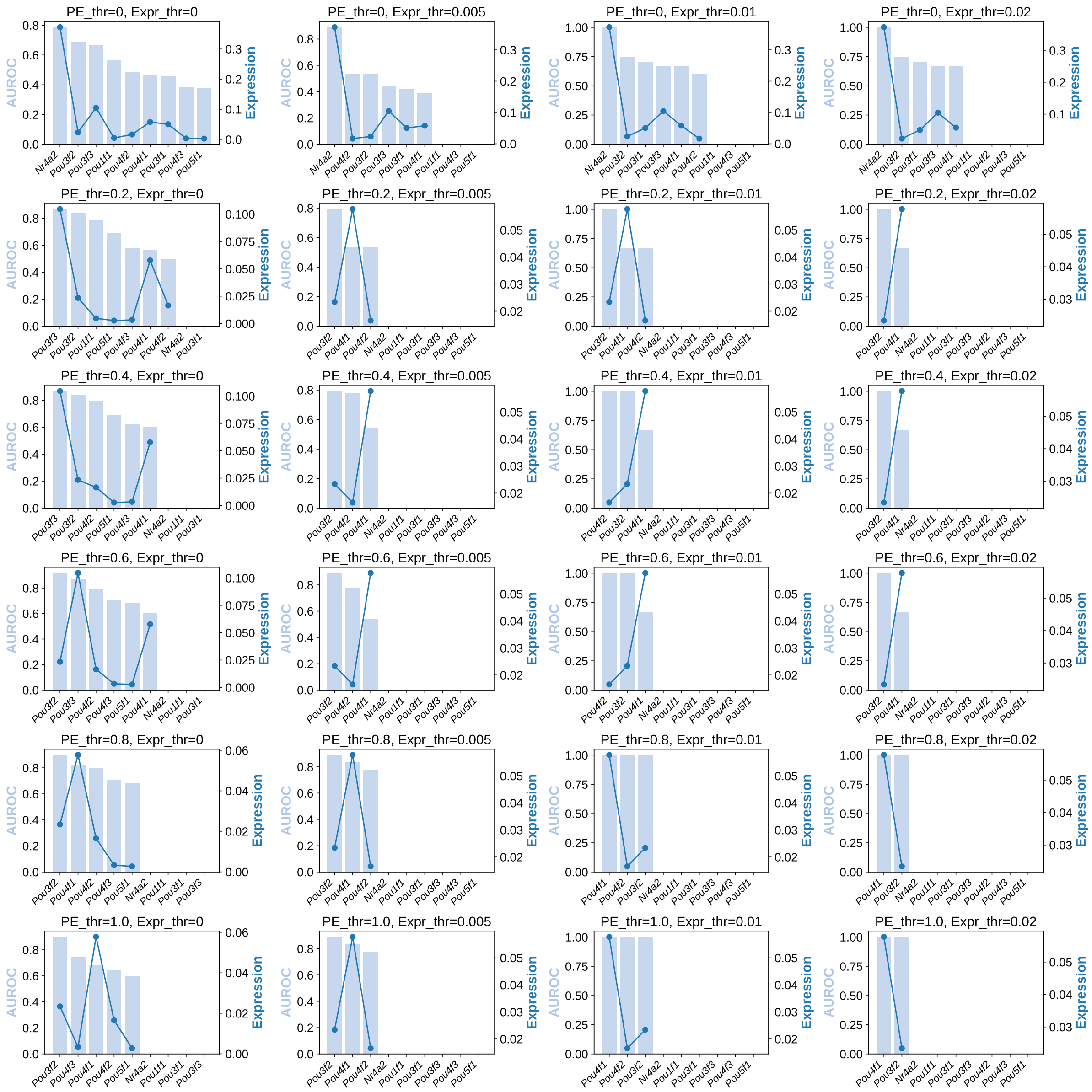

Supplementary Figure 22: (**Related to Figure 5**) Relationship between expression level and prediction accuracy for POU TFs. The dark blue line shows the expression level and the light blue bar shows the AUROC score.

#### B Supplementary Tables

Supplementary Table 1: Cell type mapping between the KO dataset and the training dataset. For one-to-many mappings, the corresponding cell types in the training dataset are separated by commas.

| Cell type in the KO data | Cell type in the training data |
|---|---|
| Arterial endothelial trajectory | Arterial endothelial cells |
| Biliary epithelial trajectory | Biliary epithelial cells |
| Cajal-Retzius cell trajectory | Cajal-Retzius cells |
| Cardiomyocyte trajectory | Atrial cardiomyocytes, Ventricular cardiomyocytes |
| Chondrocyte trajectory | Chondrocytes (Atp1a2+), Chondrocytes (Otor+) |
| Dorsal root ganglion neuron trajectory | Dorsal root ganglion neurons |
| Endocardium trajectory | Endocardial cells |
| Enteric neuron trajectory 1 | Enteric neurons |
| Enteric neuron trajectory 2 | Enteric neurons |
| Floor plate trajectory | Anterior floor plate |
| Foregut epithelial trajectory | Foregut epithelial cells |
| Germ cell trajectory | Primordial germ cells |
| Intermediate mesoderm trajectory | Anterior intermediate mesoderm, Lateral plate and intermediate mesoderm, Posterior intermediate mesoderm |
| Lens epithelial trajectory | Lens epithelial cells |
| Limb mesenchyme trajectory | Limb mesenchyme progenitors |
| Liver hepatocyte trajectory | Hepatocytes |
| Lymphatic endothelial trajectory | Lymphatic vessel endothelial cells |
| Macrophage trajectory | Border-associated macrophages, Border-associated macrophages (Cd74+) |
| Megakaryocyte trajectory | Megakaryocytes |
| Melanocyte trajectory | Melanocyte cells |
| Midgut/Hindgut epithelial trajectory | Midgut/Hindgut epithelial cells |
| Motor neuron trajectory | Cranial motor neurons, Spinal cord motor neurons |
| Myoblast trajectory | Myoblasts |
| Myotube trajectory | Myotubes |
| Neuron progenitor cell trajectory | Neural progenitor cells (Neurod1+), Neural progenitor cells (Ror1+) |
| Olfactory ensheathing cell trajectory | Olfactory ensheathing cells |
| Olfactory epithelial trajectory | Olfactory epithelial cells |
| Olfactory sensory neuron trajectory | Olfactory sensory neurons |
| Osteoblast trajectory | Pre-osteoblasts (Sp7+) |
| Pre-epidermal keratinocyte trajectory | Pre-epidermal keratinocytes |
| Primitive erythroid trajectory | Primitive erythroid cells |
| Roof plate trajectory | Anterior roof plate, Posterior roof plate |
| Satellite glia trajectory | Satellite glial cells |
| Spinal cord dorsal trajectory | Spinal cord dorsal progenitors |
| Thymic epithelial trajectory | Thymic epithelial cells |
| Venous and capillary endothelial trajectory | Venous and capillary endothelial cells |
